# Repix: reliable, reusable, versatile chronic Neuropixels implants using minimal components

**DOI:** 10.1101/2024.04.25.591118

**Authors:** Mattias Horan, Daniel Regester, Cristina Mazuski, Thomas Jahans-Price, Shanice Bailey, Emmett Thompson, Zuzanna Slonina, Viktor Plattner, Elena Menichini, Irmak Toksöz, Sandra Romero Pinto, Mark Burrell, Isabella Varsavsky, Henry WP Dalgleish, Célian Bimbard, Anna Lebedeva, Marius Bauza, Francesca Cacucci, Thomas Wills, Athena Akrami, Julija Krupic, Marcus Stephenson-Jones, Caswell Barry, Neil Burgess, John O’Keefe, Yoh Isogai

## Abstract

Neuropixels probes represent the state-of-the-art for high-yield electrophysiology in neuroscience: the simultaneous recording of hundreds of neurons is now routinely carried out in head-restrained animals. In contrast, neural recording in unrestrained animals, as well as recording and tracking neurons over days, remains challenging, though it is possible using chronic implants. A major challenge is the availability of simple methods that can be implemented with limited or no prior experience with Neuropixels probes, while achieving reliable, reusable, versatile high-density electrophysiology. Here we developed, deployed, and evaluated the real-world performance of Repix, a chronic implantation system that permits the repeated re-use of Neuropixels probes. The lightweight system allows implanted animals to express a full range of natural behaviors, including social behaviors. We show that Repix allows the recording of hundreds of neurons across many months, up to a year, with implants across cortical and subcortical brain regions. Probes can be reused repeatedly with stable yield. Repix has been used by 16 researchers in 10 laboratories to date, and we evaluated the real-world performance of Repix in a variety of chronic recording paradigms in both mice and rats with a combined 209 implantations. We found that the key advantage of Repix is robustness and simplicity. Adopters of Repix became proficient at five procedures on average, regardless of prior experience with *in vivo* electrophysiology. With the companion protocol alongside this article, the performance and user-friendliness of Repix should facilitate a wide uptake of chronic Neuropixels recordings.

## Introduction

Measuring neural activity at high temporal resolution is one of the foundational techniques in neuroscience, and methods to measure minute voltage fluctuations *in vivo* have been continuously refined for decades^1^. Recently, Neuropixels probes^2, 3^ have been transformative by massively increasing the number of neurons that can be measured electrophysiologically^4^ in mice, rats, non-human primates^5, 6^, humans^7–9^, and even octopi^10^. Notably, measurements in head-restrained animals using acute recordings benefited most immediately from this technology, unlocking new insights for the whole mammalian brain at scale^11–16^. *Chronic* implantation has two benefits over *acute* recordings: Freely moving behavioral paradigms allow animals to express a large palette of behaviors, for example foraging, decision-making, social behaviors, navigation, prey capture, and escape, and recording over days and months allows tracking of individual neurons^3^ while animals learn and update their behaviors and beliefs. Therefore, unrestrained and chronic recordings vastly expand the questions that can be addressed.

However, chronic implantations are challenging. Implantation of the Neuropixels probes must ideally balance a requirement for the stability of the implant with an option to explant the probes for later reuse, and this sets a high technical and logistical bar. For instance, initial chronic recordings with Neuropixels were conducted with the probes fully cemented in place to ensure their stability, preventing probe re-use^2, 3^. In response, retrievable systems for chronic implants have been developed. These systems aim to attach Neuropixels probes to the skull while allowing later explantation for probe reuse. Juavinett et al. developed a chronic recording in mice for Neuropixels 1.0 probes and demonstrated the ability to explant the probes^17^. In rats, a system developed by Luo et al. allowed up to four probes to be used^18^. Similarly, Van Daal et al. developed a 3D-printed dual probe fixture for use in rats, a single probe version compatible with use in mice, and a movable single probe version that allows the user to advance the probe after implantation^19^. Recently, Bimbard et al. developed a modular and lightweight approach to implanting two Neuropixels probes in mice, which was successfully implemented in multiple laboratories^20^. These systems use a common design principle consisting of docking and payload modules. This consists of permanently mounting a probe to a payload, which in turn can be secured to a docking module on the animal’s head. While the reported uses of these methods are limited for recording during solitary behaviors, these studies established stable chronic recording with Neuropixels with an option to reuse the probes upon explantation. As methods develop and mature, a central demand for dissemination is the ease with which new users can adopt the technology, though this is rarely tested or reported.

Here, we report the development, evaluation, and deployment of Repix, a minimalistic Neuropixels implant system. While prior systems have been effectively used for recording during solitary behaviors, Repix is fully compatible with a range of behaviors including social behaviors. The implant is composed of only three principal components: a cassette, posts, and a cover. Repix’s aluminum construction provides mechanical resilience for long-term implantation and social interactions. The major advantage of this design is to minimize the complexity of adoption while maintaining the stability of recording for months, up to a year. Repix can be used with any Neuropixels probe type and in both mice and rats. We shared the beta version of Repix with ten labs and further optimized it for general use. We found that Repix is readily transferable for chronic recording in a variety of applications across users and should allow rapid adoption by new users. A companion protocol details step-by-step procedures.

## Results

### The design principles of Repix

To lower the barrier of entry for chronic Neuropixels recordings, we considered multiple factors associated with the usability of chronic implants in designing our system in addition to the performance: Reliability, reproducibility, flexibility, and reusability.

#### Reliability and reproducibility

We designed a minimal system to achieve reliable and reproducible implantation of Neuropixels probes. Repix is composed of three components: a probe cassette, a pair of posts, and a probe protector cover. We used the payload and docking module principle of explantable probes similar to existing implant systems (**Fig 1A**). Each probe is permanently mounted to the cassette (payload module), which in turn slots into the posts (docking module), and the two parts are secured by multiple screws (see Supplemental Materials for CAD files). The cassette has a flat front and a convex back (**Fig 1A**, top and bottom, respectively). On the front, there is a raised edge to aid the mounting of the probe with epoxy glue. The cassette can accommodate both Neuropixels 1.0 and Neuropixels 2.0 probes. Two channels run through the length of the cassette. These are the slots for posts, as well as for the stereotaxic connector which holds the cassette during implantation and explantation (**Fig 1B** and **1C**). The posts are held in place with up to twelve M1 screws (**Fig 1A**). The posts have feet designed to sit across the skull of the animal and be enveloped in dental cement (**Fig 1A**). Both cassettes and posts are machined in aluminum (AW6082-T6) unlike 3D-printed plastics used in existing methods. While the use of aluminum increases the weight compared to a plastic alternative, the miniaturization of Repix and the simplicity of its components result in a system with a total weight of only 2.4g (cassette: 1g, posts plus 8 screws: 0.1g, Neuropixels 2.0: 0.183g, headstage: 2.0: 0.7g, headstage holder v2.0: 0.4g,). We chose to ensure maximum mechanical strength for more demanding applications, for example recording during social interactions such as rough-and-tumble fights of male mice, and in larger animals such as rats. In brief, we ensured Repix is versatile for the full range of current Neuropixels 1.0 and 2.0 probes, light enough to be implanted in mice (**Fig 1G**), and strong enough to be used in rats (**Fig 5C**).

**Figure 1.**
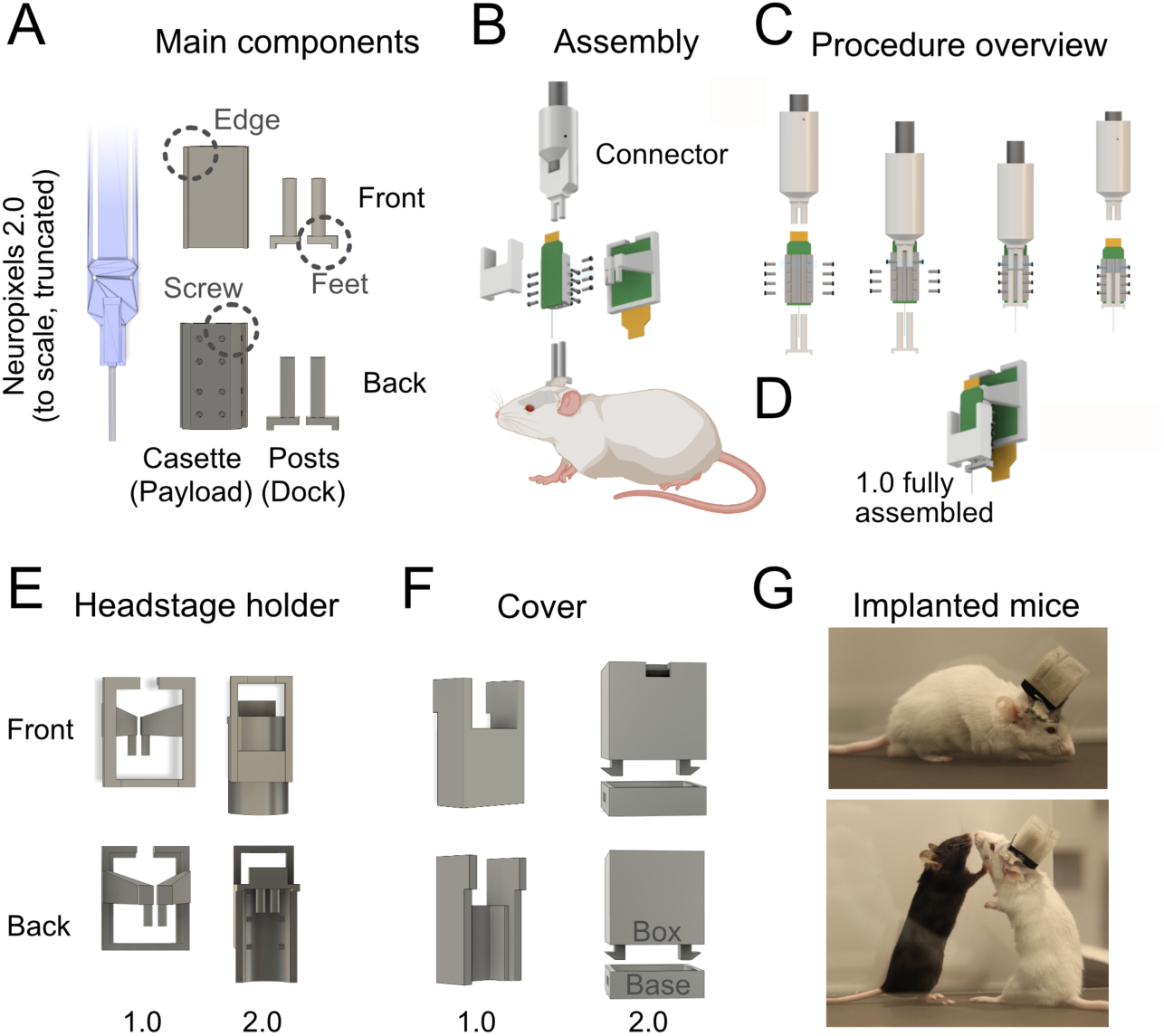
Repix is designed with few and simple components. **A.** Repix has only two principal components: payload and docking modules. Probe cassettes and posts are constructed in aluminum, shown front (top) and back (bottom). A model of a Neuropixels probe 2.0 multi-shank is shown for scale. Dashed circles highlight features that are discussed in the text (“Edge”, “Feet”, “Screw”). **B.** Chronic implantation with Neuropixels 1.0 probes. The system is assembled first by fixing the probe to the cassette. The cassette is held by a 3D-printed connector, top. The cassette and probe can then be moved independently of the posts, which lock onto the cassette with M1 screws. The cover and headstage holder complete the system. **C.** The procedure to implant the probe. The connecter is attached to the channels of the cassette and secured with screws, second panel. Posts are attached from the cranial side and secured with screws, third panel. Finally, once the probe is in place, the feet of the posts are cemented onto the skull of the animal and the connector can be loosened, fourth panel. **D.** Fully assembled implant with the Neuropixels 1.0 cover and headstage holder. **E.** The headstage can be attached with a headstage holder that fixes into the slot previously used for the connector. There are versions for Neuropixels 1.0, left, and the further miniaturized Neuropixels 2.0, right. **F.** A cover is used to further protect the shanks. The 2.0 cover consists of a base and a box that snaps into place, with access to the 4-pin connector of the headstage. CAD files for all components are available along with a detailed surgical protocol. **G.** The system in place in a mouse (CD-1, age: 10 weeks) with a 2.0 cover.

Repix combines aluminum components for strength and reliability with plastic components for flexibility. The precision-machined aluminum components minimize part-to-part variability. To ensure that the parts can be easily manufactured, multiple versions were designed and tested. This means that the aluminum cassette and posts can be reliably manufactured both in an in-house machine shop or by commercial vendors in a standardized fashion. Thereby, Repix parts can be reliably manufactured in a standardized fashion, which ensures reproducibility from procedure to procedure. We added a 3D-printed probe guard to protect the probe and other electronic components, especially the flex cable. The smaller size of Neuropixels 2.0 probe and headstage allowed further miniaturization of the system, and we implemented different guards for Neuropixels 1.0 and 2.0 probes (**Fig 1E**). The headstages for Neuropixels 2.0 are smaller than those for Neuropixels 1.0, and we developed 3D-printed headstage holders customized for each probe type. Furthermore, we designed an optional cover to fully encase the Neuropixels 2.0 implant, including the headstage, in a 3D-printed box to reliably protect from debris or cagemates damaging the implant (**Fig 1F**, right). This means users can implement the covers needed for their use cases, and the supplied files make it easy to change or update, as needed. The result is a system that is easy to manufacture, easy to implement, and reliable.

#### Flexibility and reusability

Procedurally, the implantation of a chronic recording device involves the opening of the skin, a craniotomy, sometimes a durotomy, the cementing of the system to the exposed skull, and then a closure. In all cases, the goal is to securely mount the system to the animal’s skull with the probe in the targeted brain area. Using Repix, this is achieved in a three-step process: mount the probe to the cassette, cement the posts in place, and cover the implanted probe. This is detailed in the companion protocol (https://www.protocols.io/view/protocol-for-repix-reliable-reusable-versatile-chr-davx2e7n).

First, a probe is permanently mounted to a cassette (**Fig 1C**, first panel). The cassette is then temporarily attached to a connector with screws (**Fig 1C**, second panel). The posts are put in place and also held with screws (**Fig 1C**, third panel). With the connector attached to a stereotactic frame, the probe is lowered into a prepared craniotomy. Once the probe reaches the target coordinate, there are two options: first, the surgeon has preset the post height before the surgery, and the feet can be cemented in their current position, or second, the posts are unfastened from their screws and lowered onto the skull. The screws can then be tightened again and the posts are cemented in place. The angled design of the feet of the posts allows cement to flow under and around them, ensuring a secure connection to the skull. The connector is detached and the implant is in place (**Fig 1C**, fourth panel).

For rigid probes like Neuropixels, there is a concern with tissue damage attenuating unit yield and limiting long-term stability^1^. Once the probe is in place, stability over days depends partly on limiting the amount of movement of the probe relative to the brain tissue. This is achieved by fastening the cassette to the posts with screws (**Fig 1C**), resulting in neural recording stability comparable to existing chronic implant systems with Neuropixels probes^17–20^ (see e.g., **Fig 3B** and **5C**). Although our initial procedure used twelve screws, most users opted for eight, some four. One user successfully used a single screw per post.

The last step of the implantation process is to cover the front of the implant and optionally attach the headstage. Implanting the headstage along with the probe allows a quick “plug-and-play” connection when starting an experiment and is generally recommended over attaching the headstage to the flex cable repeatedly^19^, which has been reported to cause deterioration and/or failure of the connection. However, where the implant is too heavy (e.g. for young or small animals), attaching the headstage only during the recording sessions is an option. It may simply be a preference to not implant the headstage, irrespective of the weight considerations. Eleven of fifteen Repix beta users implanted the headstage with the probe (e.g. **Fig 1D**), and the four who did not (MBa and JK, AL and CB) reported no complications or failures.

Over the last five years, the Repix system enabled users to flexibly adopt and adapt chronic recording with Neuropixels probes. Many users modified our standard protocol to optimize for their use cases. For example, the entire procedure can be completed in a single surgery, and this is done by most users. In contrast, some users have first made a craniotomy and finalized the implantation in a subsequent procedure while the animal was awake. The latter may be essential in implantations in small brain areas where physiological signatures during wakefulness can guide the targeting of the electrode. Additionally, the feet of the posts can be easily customized for optimal positioning of the implant on the skull. Some recording targets might favor angled insertions and the posts may need to be cemented in an angle. It is straightforward to modify the design of the base using the design template (Supplemental Materials). The small footprint of the posts also facilitates the use of Repix for a variety of brain areas.

A reusable chronic Neuropixels implant significantly reduces the cost of each experiment. Indeed, we achieved reusability via reversible links between payload and docking modules. Explantation is done essentially in a reverse order of implantation. First, a connector is attached to the cassette and is secured by four screws (**Fig 1C**). Next, the screws holding the posts and cassette together are loosened. Using a stereotaxic device, the cassette-probe assembly is slowly pulled out from the animal. After cleaning^2^, the probe can then be reused. Posts are also reusable. Moreover, Repix is compatible with re-implanting the probe using existing posts in the same animal.

### Repix enables reliable chronic recording and has been used by many users across multiple labs

The transferability and reproducibility across labs are essential criteria for robust methods. To this end, we shared the early versions of Repix with sixteen users across ten labs, working with both mice and rats (**Table S1**); this has already resulted in two studies published by users outside the lab that originally developed Repix^21, 22^. This allowed us to perform a real-world evaluation of Repix and assess the reproducibility of the system. To evaluate how well Repix can help implement chronic recording for new users, we reached out to all users. These researchers came with a range of surgical experience, from novice to expert. They were asked to provide feedback on each attempt associated with chronic recording in three stages: implantation, data collection, and explantation. In addition to the census on success/failure, we compiled other metadata including Neuropixels probe type, target brain area, and the presence of a grounding screw on a skull (see **Table S2**). Fifteen out of sixteen users responded. Repix was initially developed by DR, who subsequently directly trained six users in the technique (**Fig 2B**, direct training). It has also been shared without direct training from DR (**Fig 2B**, shared), and has been shared user-to-user (**Fig 2B**, black lines). This models a realistic representation of how new methods spread, and the user experiences provide valuable predictions on how easily future users will be able to adopt Repix.

**Figure 2.**
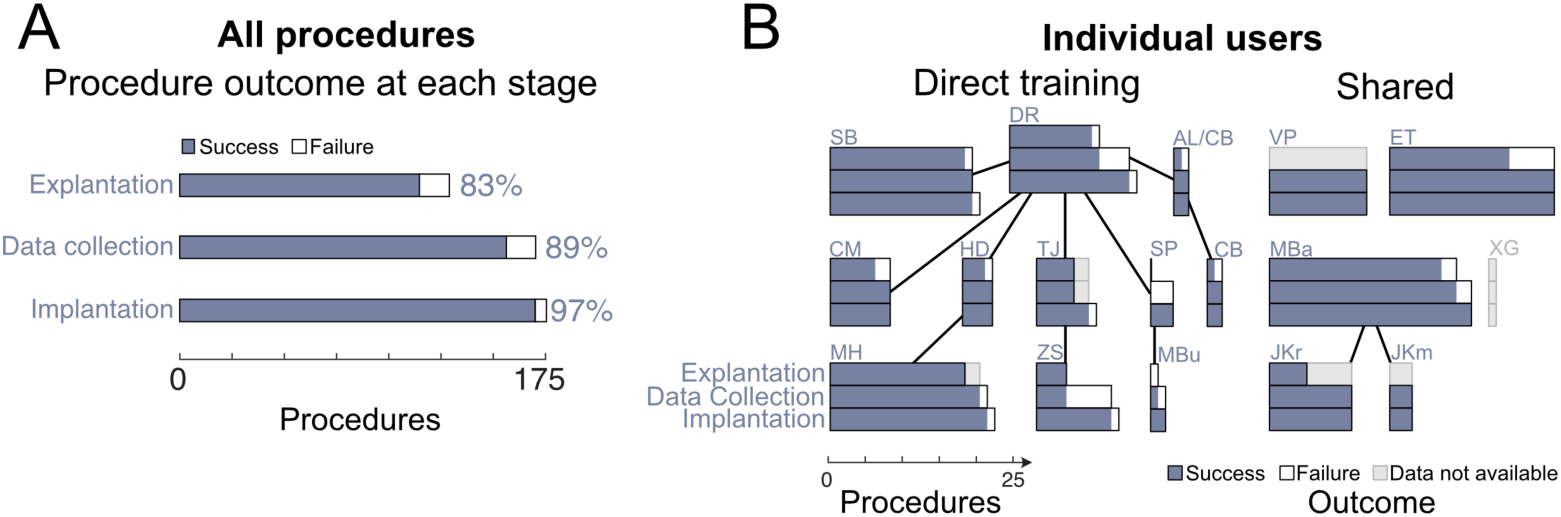
Repix has been implemented across multiple labs and users. **A.** The total number of procedures across all users in implantations, data collection, and explantation (success: purple, failure: white). These correspond to 97%, 89%, and 83% success rate, respectively. **B.** Fifteen of sixteen users responded to a survey of their use of Repix. Users had high individual success rates in implanting Neuropixels probes using Repix. The length of the bars corresponds to the number of procedures undertaken (same order as in **A**), colored for success and white for failures. Users who trained others are connected by lines. Most, but not all, users also had high individual success rates in data collection and explantation of Neuropixels probes using Repix.

Neuropixels implantations have been successfully carried out across users and labs using Repix. In total, 175 Neuropixels implantations and 127 explantations were done across all users (**Fig 2A**) with targets across the brain including striatum, amygdala, hippocampus, entorhinal cortex, visual cortex, and parietal cortex. For implantation, the mean reported number of procedures done was 10 with a mean success rate of 97% (Fig 2A, n=16). All users achieved at least 85% successful implantation. For explantation, the mean reported number of procedures done was 8 procedures with 82% success (**Fig 2B**, n=13). One user, VP, implanted probes without planning or attempting to explant them; one user, SP, had no explantation attempts; one user, JKm had no mouse explantation attempts at the time of reporting – though had successfully explanted probes in rats, indicated by JKr. There was no significant difference between the success rate when a user was in the direct training or shared groups (**Fig 2B**, direct training against shared, p=0.67).

Five failures were reported during implantation, nineteen during data collection, and twenty-one during explantation (**Fig 2A**, 97%, 88%, 83% success rates across all procedures, respectively). Typical implantation failures were probe breakage during surgery, e.g., while trying to penetrate the dura as well as accidentally breaking an exposed shank with surgical tools. Examples of failures during data collection include insufficient shielding leading to shank breakage, insufficient protection from cage mates during social housing, and implants detaching from the skull. Explantation failures included shanks stuck to silicone artificial dura or Vaseline, or inadvertent cementing of the cassette onto the posts. One project (SP and MBu) had technically successful implantations but had failures on all five of their combined procedures due to a lack of shielding. This can be attributed to the location of their target which is very anterior in the brain (frontal cortex of the mouse: AP +2.5, ML 1.5), causing the probe guard to be placed close to the animal’s eyes. One animal had an eye infection. This challenge, together with the greater curvature of the head in the anterior location, led to probe covers being placed sub-optimally, resulting in shanks breaking only a few days into data collection in four of five animals attempted (5 days in two animals, 8 days in 2 animals). All examples are incorporated in the best practices section in the protocol published along with this report. Taken together, our user experience data show that researchers with varying levels of prior surgery and *in vivo* electrophysiology experience can reliably adopt chronic Neuropixels recordings with Repix.

### Repix enables stable recording for months

To achieve chronic recordings for the same population of neurons over time, we determined if the unit yield is stable over months. Six beta users from four different laboratories measured long-term yields using Repix in mice (**Fig 3**) and rats (**Fig 5C** and **5D**). To estimate the yield and stability, we analyzed implants with three different brain targets. The included mice were assessed on both head-fixed and freely moving behaviors, including social behaviors (**Fig 3**, n=27 mice), and the implants included Neuropixels 3A, Neuropixels 1.0, and Neuropixels 2.0 probes, both single and multi-shank versions (see **Fig 5B**). Animals had implants in either hippocampus (**Fig 3B**, top row and red, n=4), entorhinal cortex (**Fig 3B**, second row and orange, n=13), amygdala (**Fig 3B**, third row and green, n=5), or V1 (**Fig 3B**, fourth row and teal, n=4). Yield was defined as the total number of units (left column) or number of good single units (middle column) recorded simultaneously using Kilosort output labels, (see Methods for details). Using Repix, we found the number of recorded units was generally high and stable across days and months, up to 364 days (**Fig 3A**). The average total unit yield was 293 (s.e.m.=29), 429 (s.e.m.=54), 440 (s.e.m.=66), 292 (s.e.m.=78) in the hippocampus (CA1 and CA3 concurrently), entorhinal cortex, amygdala, or V1 respectively (**Fig S3**, mean across mice). The average good unit yield was 95 (s.e.m.=35), 159 (s.e.m.=21), 94 (s.e.m.=22), and 96 (s.e.m.=26) respectively. These data were collected during a range of behaviors, including foraging, virtual navigation tasks, and social interactions, lasting at least 15 minutes and up to 90 minutes per recording. Neuropixels probes allow users to select up to 384 recording channels from the 960, 1280, or 5120 available electrodes (for Neuropixels 1.0, 2.0 single shank, or 2.0 multi shank, respectively). For each animal and insertion, custom maps of selected electrodes were created by the user. Typically, the unit map was stable across days, though at the discretion of the user, it could be updated to accommodate experimental needs.

**Figure 3.**
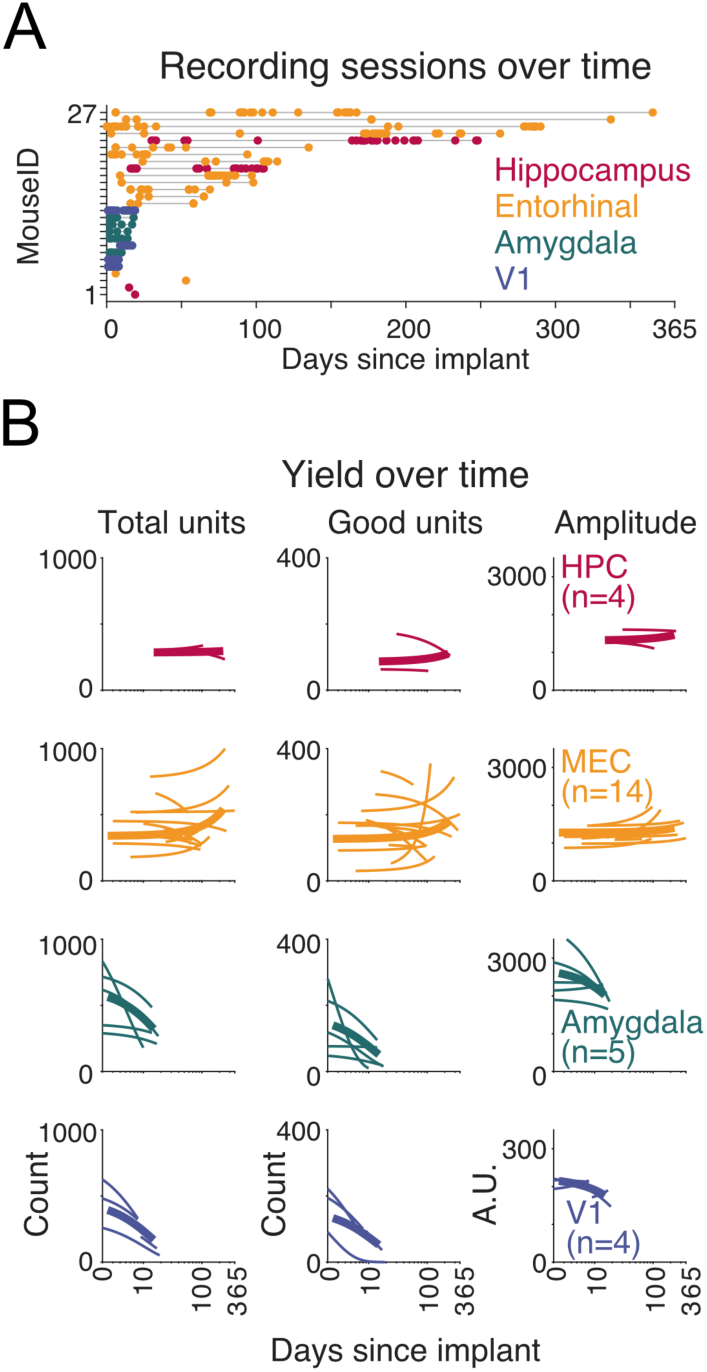
Repix can be implanted for up to a year with stable unit yield. **A.** The summary of recording sessions in 27 mice implanted with Repix. These implantations were in the hippocampus (red), entorhinal cortex (orange), amygdala (green), and V1 (blue). The maximum duration was 364 days. **B.** Individual implants maintained stable unit yield over weeks and months, both in total units (left) and good units (middle). These units had a stable waveform amplitude over time (right). Thin lines are logarithmic fits to the recording data from the individual and thick lines represent logarithmic fit to binned average across animals. Note the logarithmic time scale.

In most of these mice, we did not observe an initial decrease in yield over the first seven days (**Fig 3B**). That is, Repix recordings behaved similarly to a report by Bimbard and colleagues in mice^20^ where initial attenuation was limited (or absent), as opposed to the systematic unit decrease observed by Luo and colleagues in rats^18^. However, we observed a substantial attenuation of yield in the amygdala and V1. The mean change in good units was −0.0016 units/day (s.e.m.=0), 0 units/day (s.e.m.=0.0023), −0.13 units/day (s.e.m.=0.07) and −0.22 units/day (s.e.m.=0.09) in the hippocampus, entorhinal, amygdala, and V1 respectively (**Fig S3**). Taken together, these data suggest that Neuropixels recordings can be successfully undertaken over months using Repix with yield and stability in line with or better than other retrievable implant systems.

### Repix allows the reuse of probes

While Repix allows the reuse of probes, not all users opted for this capability. Nine of fifteen users supplied data on reusing probes. Probe reuse generally increased as users gained proficiency and started to produce data consistently. Across respondents with reuses, 30% (36/122) of all implantations were with reused probes (**Fig 4A** and **4B**). The maximum number of reuses for single probes was 5 times (**Fig 4C**), conducted by SB. These experiments targeted the amygdala where the unit attenuation was significant within a week of implantation therefore the implant-explant cycles were shorter. To assess the impact of multiple uses of an individual probe, we focused on the yield on the first recording day. Across nineteen implantations in the amygdala, the initial yield for a probe did not systematically differ with the number of reuses it had undergone (**Fig 4C**, 2-way ANOVA, F=0.09, p=0.98). There was no systematic change from early to later implantations done by user SB. Equally, there was no difference in yield by the individual probe used (**Fig 4C**, individual probes, ANOVA, F=0.68, p=0.65).

**Figure 4.**
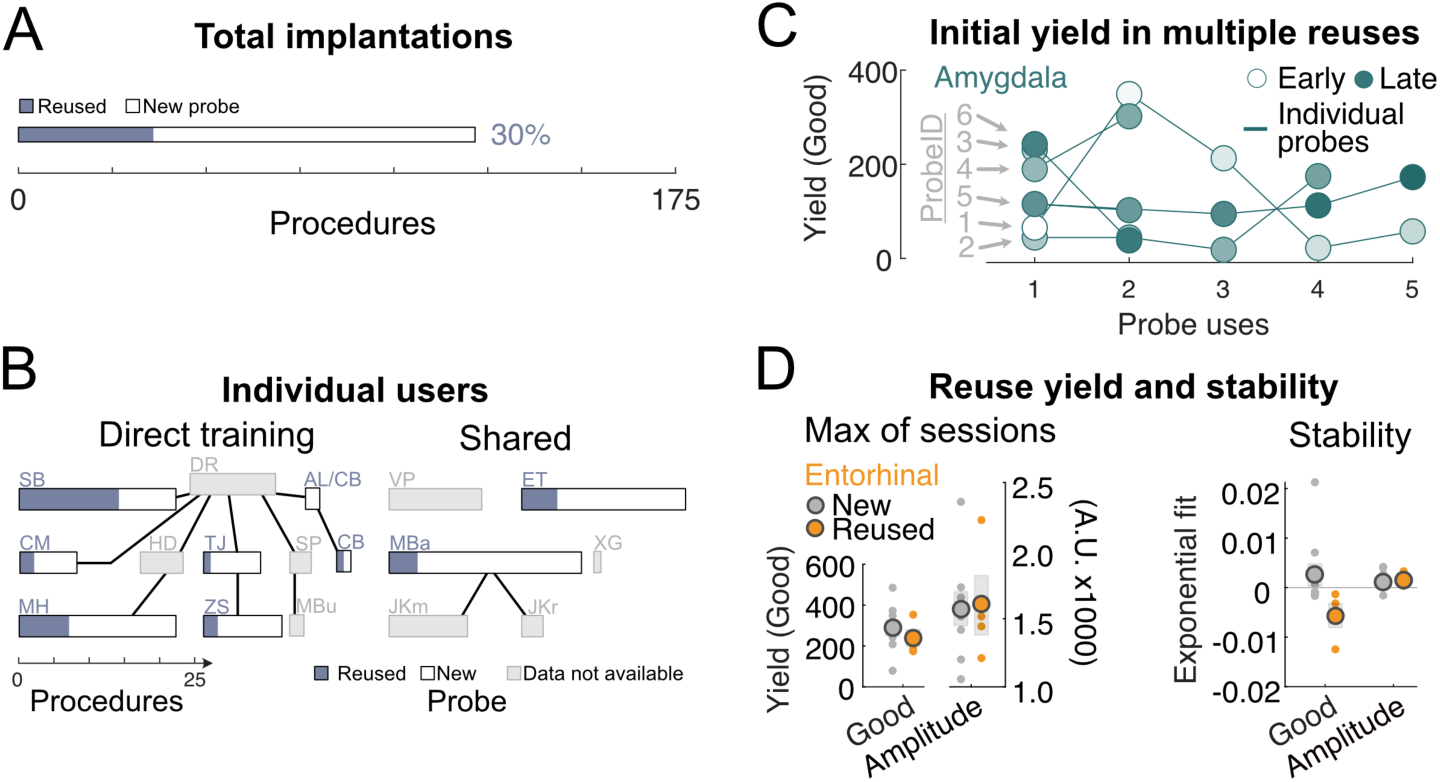
Repix allows reuse of Neuropixels probes. **A.** Across all users, 30% of implantations were done with reused probes. **B.** Nine of fifteen users opted to use Repix’s feature to reuse probes. Seven of fourteen users supplied data on reusing their probes. Colored bars indicate the number of implantations undertaken with a reused probe. **C.** The initial-day yield of good units in the amygdala did not decline as the same probe was reused multiple times. Data points from individual probes are connected by lines. Circles are colored by their surgical date rank, early to late implants from light to dark. **D. Left:** New (gray) and reused (orange) probes implanted in the entorhinal cortex had similar unit yield, defined as the peak number of units in a single session, left axis. Equally, waveform amplitudes were similar, right axis. The dots represent data from individual implants. The mean of implantations is represented in the circles with s.e.m. in the shading. **Right:** Exponential fits show no significant difference in good unit yield stability. Waveform amplitude was similarly stable. Same color conventions as in left panel. A significantly quicker drop-off in total units was observed (**Fig S3**).

To assess the yield and stability for longer recordings, we next turned to the implants in the entorhinal cortex. Here, four probes were reused and eleven were new. Reused probes had similar yield as measured by the highest number of units in a single session (**Fig 4D**, left panel, leftward axis, n=4 reused probes in orange compared to 11 unused probes in gray, all in entorhinal cortex), and as measured by the average yield across all sessions (not shown). Further, average amplitudes for spikes collected with probes that were reused were not significantly different than those collected from new probes (**Fig 4D**, left panel, rightward axis). The fitted exponential decay was not significantly different for good units or amplitude (**Fig 4D**, right panel), suggesting similar decay characteristics in yield and waveform amplitude between reused and new probes (p>0.05 for all comparisons in **Fig 4D**). The only difference was in the stability of MUA units (**Fig S4**, p=0.0014), in which there was a larger multi-unit decay in the implants with reused probes in this small subsample. Taken together, Repix enables long-term recording with good yield in several brain regions. We show that the reuse of probes could result in a minor decrease in the recording quality, but this does not compromise the reuse from a practical standpoint.

### Repix can be used across different experimental paradigms

To evaluate the capability of Repix to be used for a wide range of use cases, we tested how the weight and the height of the implant affected the unrestrained movement of mice during social interactions. We used an inertial measurement unit (IMU) to record the angular velocities of pitch and roll of the animals’ heads with and without a Repix implant during bouts of interaction with males, females, and pups (see Methods). Comparing Repix+IMU mice to IMU-only mice, movement dynamics were similar (**Fig 5A**). In fact, implanted animals moved similarly to their IMU-only counterparts across high and low movement frequencies. Average pitch and roll velocities were not significantly different (pitch p=0.11; inset, roll=0.39). As part of deployment across labs, Repix has now been used in mice in head-fixed virtual reality, spatial navigation in mazes, and social interactions, and has been well-tolerated by the mice in all these contexts.

**Figure 5.**
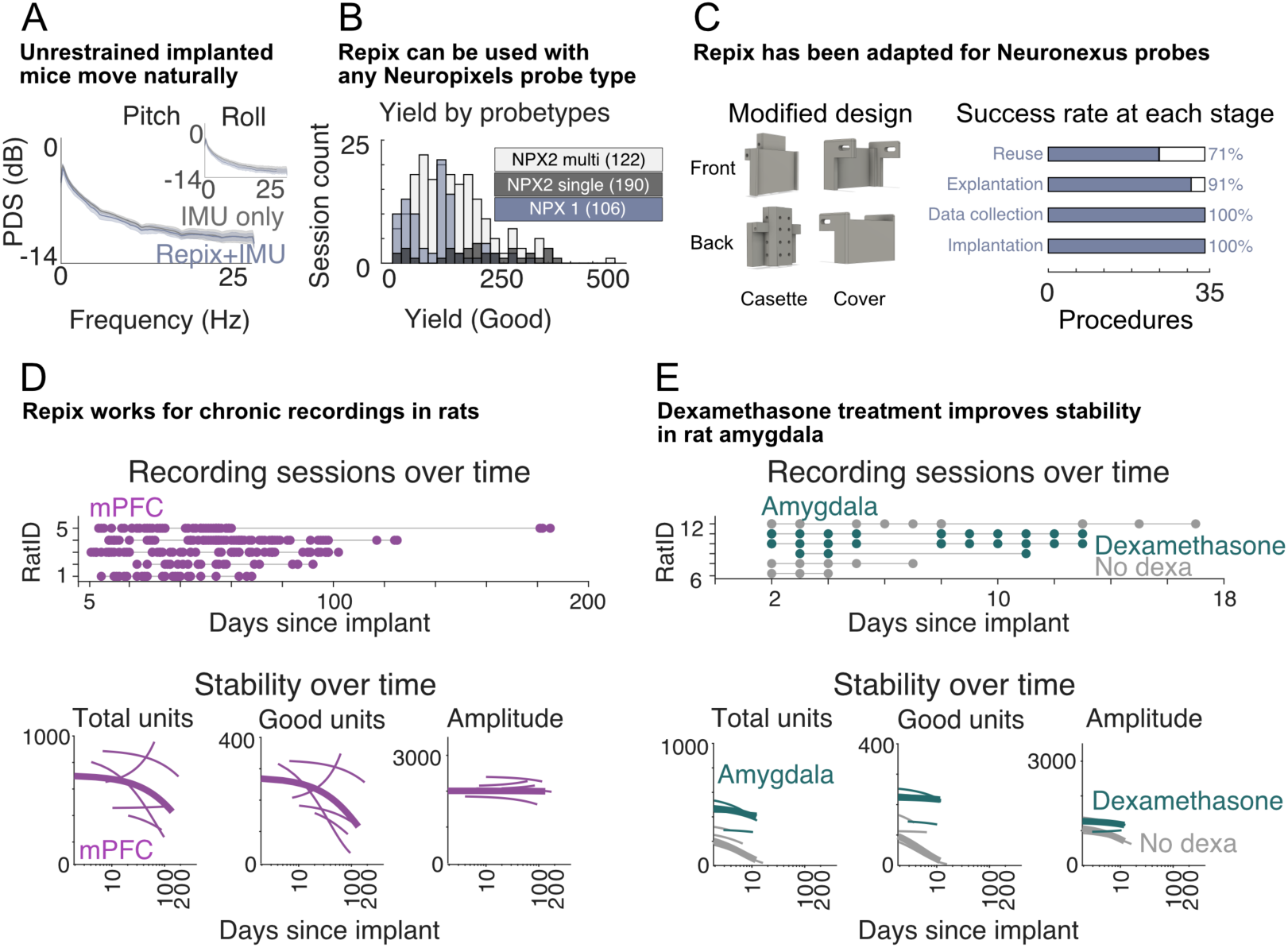
Repix can be used in many different experimental paradigms. **A.** Unrestrained implanted mice move naturally. Implanted mice (purple) showed similar movement dynamics to non-implanted mice (gray) across the spectrum of slow (low frequency) to fast (high frequency) head movements during social interactions, both in pitch (main panel) and roll (inset). **B.** Repix can be used with any Neuropixels probe type. Histogram of good unit yields by probe type, Neuropixels 1.0 in purple, Neuropixels 2.0 single shank in dark gray, Neuropixels 2.0 multi-shank in light gray (mean yield in legend). While the users reported good yields across areas and probes, each user tended to use a single probe type in each brain area, making direct comparisons of yield by type within an area difficult. **C. Left:** Repix parts for Neuronexus probes. **Right:** Success rates of each experiment stage, including reuse of probes, using the modified design. **D. and E.** Repix is fully compatible with chronic recording in rats. Stability and yield from mPFC/ACC (D, n=5 rats) and amygdala recordings (E, n=6 rats). Same conventions as Fig 3. **E.** Rats injected with dexamethasone (green, n=3) had improved yield and stability compared to non-injected animals (gray, n=3).

Through development and deployment, Repix has been used with all types of Neuropixels probes (all commercially available 1.0 and 2.0 probes, single and multi-shank, as well as pre-commercial Neuropixels 3A and 2.0 probes). Most experiments were conducted using single types. While we do not have a sufficient number of targets of the same area with different probes to test the effect of probe type on yield or stability within a targeted brain area, users reported high average yields with any of the probe types (**Fig 5B**, mean yield by session is 106, 190, 122 for Neuropixels 1.0, 2.0 single shank, and 2.0 multishank, respectively). Repix can even be modified to accommodate other silicon probes including Neuronexus probes (file in Supplemental Materials). One user (IV) widened the cassette and cover but used the identical protocol (**Fig 5C**, left, and see Methods). Across 34 implantations, IV achieved 100%, 100%, and 90% success rates for implantations, data collection, and explantation, respectively (**Fig 5C**, right). Reuse was also possible with this modified system, and 71% of implantations were done with previously used probes (**Fig 5C**, right). This use case exemplifies that the supplied CAD files can be further modified to accommodate future needs for different probe types. This design is also added to the repository. In conclusion, these use cases suggest that Repix is a versatile platform allowing customized solutions for chronic recording.

We next evaluated Repix implants in rats. Four of fifteen users applied Repix in rats (see users CM, VP, MB, JKr in **Fig 2B**. In addition, user IV applied the Neuronexus-modified Repix in rats, too, **Fig 5C**). While the weight of the implant is less of a concern for rats, the protection of the implant is paramount, because rats are larger and stronger (see Methods and Discussion for rat-specific surgery practices). Repix provides sufficient mechanical resilience for use in rats, and the rats tolerate the implants well. Two users employed dual-probe implants in rats (see Methods). Repix has been used in rats doing a plethora of unrestrained behavioral tasks, including social behavior^22^, foraging, object exploration, working memory, and operant tasks. Implanted rats displayed typical social behavior including social sniffing, allogrooming, chasing, and mating^22^. In addition, two users made dual-probe implants (see Methods). Next, to assess the yield and stability of these implants, we analyzed recordings from the two users from two different labs, targeting the medial prefrontal cortex (mPFC) and amygdala, respectively. In the rat mPFC, the yield was high and stable for more than 100 days (**Fig 5D**, n=5 rats), with recordings beyond 100 days in two animals. The average total unit yield was 578 (s.e.m.=99) and the average good unit yield was 196 (s.e.m.=38) in the mPFC (**Fig 5D**, purple, n=5). The mean decrease in all units across rats was −0.0027 units/day (s.e.m.=0.0031), and in good units it was −0.0056 units/day (0.0049) (**Fig 5D**, gray fit line, n=5). In the rat amygdala yield was generally high, with decay constants comparable to those in the mouse amygdala recordings (**Fig 5E**, n=6 rats). The average total unit yield was 211 (s.e.m=56) and the average good unit yield was 102 (s.e.m.=32) (**Fig 5E**, gray, n=3). The mean decrease in all units across rats was −0.081 units/day (s.e.m.=0.026), and in good units it was −0.095 units/day (s.e.m=0.056) (**Fig 5E**, gray fit line, n=3). We also analyzed recording data in the amygdala, in which animals were treated with dexamethasone, a synthetic glucocorticoid used to decrease inflammation. The average total unit yield was 394 (s.e.m.=54) and the average good unit yield was 200 (s.e.m.=30) in the amygdala of rats treated with dexamethasone (n=3). The mean decrease in all units over time was −0.015 (s.e.m.=0.01), and in good units it was −0.011 (s.e.m.=0.02) (**Fig 5E**, dexamethasone cohort in orange, n=3). The decay of unit numbers was slower (non-dexamethasone group vs dexamethasone group: max yield all units p = 0.047; max yield good units p = 0.023), more similar to recordings in the hippocampus and entorhinal cortex of the mouse. Therefore, peri-operative dexamethasone appears to provide improvements in unit yield over days. Taken together, these data demonstrate that Repix enables robust and stable yields in multiple brain areas during chronic recording in rats.

### Repix lowers the barrier of expertise transmission across labs

A major hurdle in adopting a newly published technique is the steep learning curve for new users, particularly if they are external to the group that developed the technique. We therefore decided to analyze learning curves for our Repix users. We collected success and failure data from individual implantation attempts from nine out of fifteen respondents (**Fig 6**, top). All attempted implantations were categorized into one of the following categories: practice, implantation failed, data collection, explantation failed, and full procedure. Failed implantations were cases when a probe was damaged and became unusable. Collected data include animals in which the users successfully implanted probes and collected electrophysiological data, but subsequently had a probe failure, e.g., due to insufficient protection. We defined proficiency as three sequential successful full procedures, and the estimated learning curve is the number of procedures to reach proficiency. The median number of procedures done to achieve this was five (**Fig 6**, bottom, n=9, interquartile range = 3-9). This number covers a breadth of real-world experiences: Users of Repix spanned a variety of experimental and surgical backgrounds, from students with no prior surgical experience to postdoctoral researchers with more than a decade of experience. Some targets may be surgically more challenging (e.g. lateral or posterior approaches). Even with differences in background and experimental needs, all respondents became proficient with the technique in the follow-up period.

**Figure 6.**
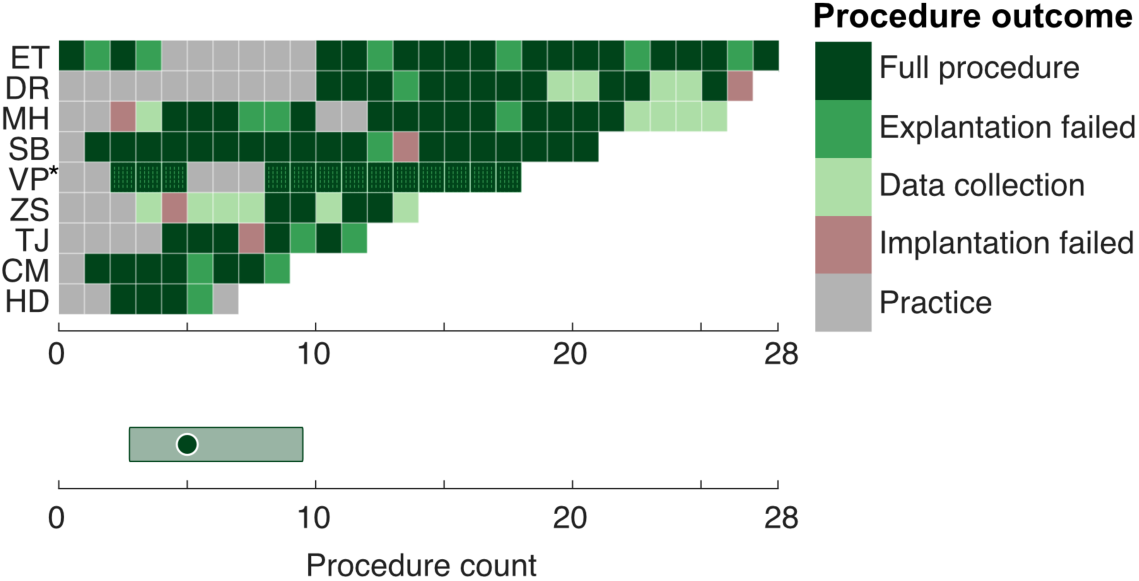
Users quickly gain proficiency. **Top:** Each procedure (attempted/planned implant) was categorized as either: Practice (gray), Implantation failed (red), Data Collection (light green), Explantation failed (green) or Full procedure (dark green), for nine users who supplied detailed surgical records. *User VP implanted Repix without intention to explant, and therefore Full Procedure includes only Implantation and Data Collection (Dark green, dotted). **Bottom:** Proficiency was defined as three sequential successful Full procedures, and the median number of procedures to reach this point is plotted (green circle, IQR in shading).

## Discussion

### Repix, a minimalistic implant system for Neuropixels probes

In this study, we developed, deployed, and evaluated Repix, a robust and reusable Neuropixels implantation system. As the Neuropixels recording technology evolved and matured, researchers developed solutions to chronically implanting Neuropixels probes. They have a common design principle using payload and docking modules and have been effectively used for some combination of recording from Neuropixels 1.0; Neuropixels 2.0; in mice or rats; with the ability to reuse probes^3, 17, 19, 20, 23–26^. Repix combines all these qualities to ensure long-term recordings in an aluminum system that is strong enough for rats and light enough for mice; which can accommodate any probe type; in assays that span rodent behaviors, including freely-moving, head-fixed, and social behaviors; where probes can be reused; all in a solution with a minimal number of components that can be adapted for individual use cases. It has been assessed through the largest number of procedures (175 Neuropixels implantations and 34 Neuronexus implantations across 10 labs) of any reported system, and we presented the real-world experience of Repix users, both advantages and disadvantages, to help guide and set expectations for future users. While all published methods have detailed protocols associated with their publications, it is often unclear to a beginner how straightforward it is to adopt a new method since the information about how well the techniques have been recapitulated outside of developers’ laboratories is often not included (for examples of systems used beyond the lab in which they were developed, see ^3, 20^). Therefore, our learning curve data for successfully implementing chronic Neuropixels implantations should provide a realistic estimate of future adoption.

For Repix, we prioritized ease-of-adoption as we consider it to be a major factor in lowering the barrier to entry. A similar approach has been taken by the Neuropixels community, which has been successful in promptly developing methods for data recording (Open Ephys^27^, SpikeGlx), processing (CatGT), spike sorting and curation (Kilosort^28^, phy, SpikeInterface^29^), analysis (NeuroPyxels^30^, BombCell^31^, PinPoint^32^, UnitMatch^33^). The grass-roots community of support and detailed documentation have proven useful in supplementing (or even replacing) the traditional model of teacher-tutor relationships in adopting challenging techniques. We therefore implemented an open-source solution that can be easily communicated and implemented externally as a major feature of this system, and this report along with the CAD files and accompanying step-by-step protocol achieves this.

Notably, we designed Repix as an implant system to record neural activities during social interactions of mice. The recording in social contexts requires multiple unique considerations. First, the implant must be resilient to dynamic social interactions that often accompany fast and forceful movements, for example, during fighting. Second, to facilitate the natural social behaviors of the recorded mice, the implant must be lightweight to minimally perturb head motions. Third, the interaction partners have to be able to naturally interact with the recorded mice, without causing entanglement with the implant parts and wiring. To meet these requirements, we decided to use aluminum machined parts, instead of 3D-printed plastic, for the core components of the implant system. The enclosures for the probe cassette and headstage provide additional protection for the probe and cables during long-term social housing (**Fig 1F**), as cagemates chewed the wires in early iterations of the design. To help ease the adoption of the methods, we iteratively designed the parts so that they can be manufactured in a standardized fashion and at a low cost (∼30 GBP per implant in 2020 from 3D Hubs, comparable in 2024). Yet, some beta users reported higher costs of aluminum parts in the US (e.g., ∼100 USD per implant from Xometry). This could likely be mitigated by sourcing from the vendors that will allow international shipping, or by making bulk orders. The same users reported that outsourcing added complexity and time to iterate potential design changes. We made a comprehensive materials and tools list to help estimate the cost for the full Repix system and associated materials and tools (Supplemental Materials, as well as in the protocol).

A solid and simple design of Repix allowed us to extend this platform across different recording sites; for different behavioral assays; and for recording in rats. To ensure that the method is reproducible among several researchers, we have shared the method with more than a dozen scientists across ten laboratories. After approximately five years of testing, we are confident that the Repix system is suitable for general use.

### Repix allows stable long-term recording and reuse of probes

An effective chronically implanted probe should allow stable recording over time from hundreds of single neurons in unrestrained animals that express a diverse behavioral repertoire. Here, we show that Repix can indeed be used for experiments running across weeks and months, with the longest recording duration lasting 364 days. We found that attenuation in entorhinal cortex and hippocampus was low, similar to that reported by Bimbard and colleagues^20^ where initial attenuation was limited (or entirely absent). In fact, we observed that yield trajectories varied, i.e., were increasing, stable, and decreasing with time (**Fig 3B**, **4D** and **4E**), similar to findings by Bimbard and collegues^20^. This is in contrast to the systematic increase in unit decay along the anterior-posterior as well as ventral-dorsal axis that has been reported for recording in rats^18^. Users recording from the amygdala indeed found substantial attenuation. Importantly, this stability problem was greatly alleviated with the use of peri-operative dexamethasone in rats. Furthermore, no user reported any issues indicative of deteriorating mechanical stability of the implant over time (e.g. more failures as experiments progressed in time). Taken together, Repix is appropriate for long-term recordings, months up to a year, in line with the best reports from other methods.

The reuse of Neuropixels probes significantly lowers the cost of chronic Neuropixels experiments. In order to reuse probes, both implantation and explantation must be successfully achieved. And across users, explantation was high at 83%. This is similar to the Apollo implant, which is 86% when taking into account all probe errors, and only slightly lower than the 10/11 successful retrievals in the single lab experience in rats in Ghestem et al^26^. Repix has been used more times (e.g. 175 Neuropixels implantations to 31 in Apollo), across more labs, and while this might be expected to lead to more diversity of outcomes, success rates remained high. Regarding probe reuse, previous studies have included thorough testing of noise levels after initial implantation and found slight increases in noise, but within the level that made the reuse of probes appropriate^18^. In our cohort, reuse rates were not generally very high: 30% of implantations were done with reused probes. We found that users preferred to use new probes whenever possible and only reused probes later in their experimental journey. The total number of reuses of single probes (5) is comparable to the best reported^20^, and we found no meaningful detriment to the recording quality in repeated reuse.

Finally, we show that Repix is appropriate for use across all available Neuropixels probes and even Neuronexus probes. Comparing the yield and stability of different probe types in the same brain area is outside the scope of this study, as users tended to use different probes for different targets, and any differences in yield or stability between probes are highly correlated with the targeted area.

### Drawbacks and future directions

Currently, the major drawbacks of Repix are threefold. First, at 2.4g, Repix is 0.3g heavier than the lightest implant system that uses 3D-printed plastic parts. For comparison, the Apollo system weighs 1.4g without the headstage, which adds 0.7g for a total of 2.1g (the Apollo system weighs 1.6g for Neuropixels 1.0, total 2.3g). This makes the adoption in small mice, for example, under 20g, challenging, particularly if the user intends to also use a headplate. One remedy for this drawback is to use mouse strains that have an overall higher weight. For our work, we mainly used C57BL6/J animals but we also used CD-1 animals which are larger and can tolerate heavier implants for the study of social behaviors. While the use of standardized inbred strains is critical for reproducible science, one could look beyond C57BL6/J mice, a standard strain of choice by many neuroscientists for mouse behaviors^34^. A second possibility is to implant the probe without its headstage and only add the additional weight counterbalanced during tethering, a procedure used successfully by MBa and JK in freely moving animals and AL and CB in headfixed animals, or as part of the implant once the animal has grown.

The second drawback is that the current Repix design does not allow two probes to be simultaneously inserted using the same posts. Dual probe implants in mice are the standard feature of Apollo or similar systems^19, 20, 24^, and a recent report showed a 3.4g fixture with 6 probes chronically implanted^25^. The resources accompanying this article include two versions of dual-probe Repix prototypes (Supplemental Material: CAD/Cassette/Prototypes/DualProbe.step). In addition, multiple labs have implanted two Repix modules in the same rats.

Third, two users attempted to implement Repix but ultimately chose to use an alternative system (SP and MBu). Their failures prompt a need for refinement if a very anterior brain area is targeted. Additionally, two users (AL and CB) initially used Repix^21^, but decided to develop an alternative system for a lighter system that can accommodate two probes in mice (Apollo^20^). Best practices are summarized in the companion protocol, which includes lessons learned from these cases.

A prominent future direction of next-generation implant development is to incorporate features necessary for chronic implantation into the probe itself, for example, to incorporate features that will enable simple attachments to a docking module. This might also require further miniaturization, and perhaps a standardized rig for probe implantation and explantation. A concerted advance in the probe and implant designs will likely provide a refined approach to chronic recording using Neuropixels probes.

## Methods

### Subjects

In the UK, experimental procedures were conducted according to the UK Animals Scientific Procedures Act (1986) and under personal and project licenses released by the Home Office following appropriate ethics review. In the US, all procedures were performed in accordance with the National Institutes of Health Guide for the Care and Use of Laboratory Animals and approved by the Harvard Animal Care and Use Committee.

A total of 136 mice and rats were used in this study. Of them, electrophysiology is included for 40 animals. Male mice of strains C57BL6/J or CD1 (Charles River) were implanted from 12 weeks of age and at least 27g at the time of surgery unless otherwise specified. One user (ZS) implanted mice at age 9 weeks. Male rats (Lister Hooded from Charles River) were implanted from 12 weeks of age and at least 350g at the time of surgery.

### Repix

All components of Repix were designed in Autodesk Fusion 360, and machined in aluminum (AW6082-T6), either in an internal facility at the Sainsbury Wellcome Centre or by an external company (3D Hubs). M1x3mm screws were used (Yahata Neji Cross Pan, RS-online). The outer dimensions of each component are as follows. Cassette: 15mm x 8.8mm x 4mm (plus 1mm with the raised edge) with 2.1 mm diameter channels running through and four rows of M1 tapped screw holes. Posts: 2mm diameter, either 9mm or 12mm length with 4.2mm feet with an additional 0.7mm toe. CAD files for each part are attached to this article.

### Neuropixels probe mounting

Throughout the study, we used Neuropixels 1.0, including the pre-release version (3A) and commercially available version (3B), or Neuropixels 2.0, including pre-release single and multi-shank probes. The details and step-by-step of the mounting procedure, along with implantation and explantation, can be found in the protocol associated with this article. The cassette is held with the connector in a horizontal configuration. All screws are added. The probe is placed on the flat front of the cassette and held with Blue Tack (Bostik). Graph paper was used to ensure the probe to be in parallel with the connector. After verifying the alignment, the probe is then secured with epoxy glue (e.g., Araldite). The glue is allowed to fully cure, usually overnight. Posts are added and fixed with screws. Most users opt to short the ground and the reference at this point with a silver or stainless steel wire. In addition, to visualize the probe tracks after explantation, the shanks can be coated with fluorescent dye (e.g., DiI solution, ThermoFisher).

### Implantation

Repix is compatible with implanting in either awake or anesthetized animals. A standard implantation procedure is described with the protocol associated with this article. Each beta tester was sent an initial version of the protocol, and each user customized their procedures according to the requirement of experimental systems (e.g., mouse vs rat, whether or not recording needs to be made while the probe is inserted). Most users did practice surgeries usually with dummy (inert) probes on freshly euthanized cadaver animals, as allowed by local animal licenses (gray entries. CM did practice surgeries with pulled pipettes as probe stand-ins). Relevant deviations are incorporated into the standard protocol for completeness and clarity. Here, we describe the specifics of the surgery by the labs, as they differ slightly:

#### Akrami

Male Lister Hooded rats were used in this study (n=18). Animals were obtained from Charles River Laboratories and housed under standard laboratory conditions with a 12-hour light-dark cycle. All experiments were performed in accordance with UK Home Office regulations (Animal Welfare Act of 2006) following local ethical approval.

Before surgery, rats were habituated to handling and the experimental environment. Rats were administered the following drugs subcutaneously: Vetergesic: 0.003-0.017 ml/100g; Metacam: 0.02-0.02 ml/100g; Dexadreson: 0.05 ml/100g; Marcaine: 0.2 ml/100g.

General anaesthesia was induced and maintained using isoflurane (5% and 1.5%-2% respectively) delivered via an anaesthesia mask. The head was shaved, and the eyes were lubricated with eye gel to prevent them from drying. Body temperature was maintained at 37°C using a heating pad throughout the procedure.

Using a stereotaxic frame (Kopf Instruments), the skull was exposed and cleaned. Craniotomies were performed over the target brain regions at specific coordinates relative to bregma: AP 3.2mm, ML 0.7mm, targeting medial prefrontal cortex (n=7); AP −3.72mm, ML 2.5mm, targeting posterior parietal cortex (n=7); AP −3.6mm, ML 6.5mm, targeting auditory cortex (n=3).

Prior to insertion, the probe was coated with DiI (1 mM in isopropanol, Invitrogen). After craniotomy, dental cement (C&B SuperBond) was meticulously applied to the skull to augment the stability of the implant. Following the removal of the dura, Neuropixels probes were carefully inserted to the desired depth using a motorized micromanipulator (Sensapex, uMp-4) at speeds ranging from 2-15 um/second. Throughout the procedure, the brain surface was consistently moistened with sterile saline to prevent desiccation.

Once the probe reached the desired position, the metal posts of the implant were secured using UV-curable cement (3M, RelyX Unicem 2) after the removal of the micromanipulator arm. To safeguard the probe and the electrical components of the implant, the surgical site was sealed with a removable silicone sealant (Kwik-Sil sealant, World Precision Instruments). Additional application of UV-curable cement was performed to reinforce the probe’s stability and provide mechanical support for the implant.

Furthermore, to enhance protection and stability, the implant was carefully wrapped with Coban band (3M). This additional layer ensured added security and minimized the risk of damage post-surgery.

Rats were allowed to recover in individual cages until they reached their pre-surgical weight and a minimum of 48 hours. Post-operative analgesia was provided via oral administration of metacam tablets.

#### Barry

Pre-surgery: mice were habituated to handling over a 5-day period, as well as pre-exposed to non-medicated jelly in their home cage for 3 days. On surgery day: each animal was put under general anaesthesia in an induction chamber (5% Isofluorane). The following medication was administered: Rimadyl (5mg/kg), subcutaneous injection; Dexamethasone (5mg/kg), intraperitoneal injection. The animal’s head was shaved, and the animal was placed in a stereotaxic frame and on a heated pad. Anaesthesia was reduced and kept at 1-2.5% Isofluorane for the duration of the surgery. The animal’s eyes were lubricated and covered to protect them from light. The shaved surface of the animal’s head was sterilised with hibiscrub. The animal’s body temperature and breathing were monitored throughout the surgery.

Once the level of anaesthesia was satisfactory (confirmed by the absence of pedal reflex), a small incision was made at the midline of the animal’s skull. The skull was exposed and scored with a scalpel. Once the skull was levelled, the coordinates for craniotomy as well as two contralateral locations for skull-screws were marked on the skull. Two screws, one with a grounding pin attached to it, were inserted. The skull was briefly dried using cotton buds to ensure better adhesion and a protective well was formed around the location of the craniotomy using fast-curing RelyX resin cement (3M). A custom-made head fixation post was placed anteriorly to the bregma and fixed in position using dental cement (C&B SuperBond). A 1 mm craniotomy was made inside the protective well using a biopsy punch (KAI Medical) over the target brain regions at specific coordinates relative to bregma: AP 3mm, ML 2.4mm, DV 3.7mm targeting V1 and CA1 (in seven animal AP coordinate 2.7mm was used). Care was taken not to damage the dura. The craniotomy was immediately covered with Dura-Gel (Cambridge NeuroTech), which was left to cure for 15-30 mins. Then, a Neuropixels probe (sharpened and dipped in Dil) was lowered using a programmed micromanipulator (Sensapex) set to 2 µm/s. Once the desired depth was reached, the posts secured to the cassette were dropped and cemented to the skull/headpost. The probe was connected to a ground wire to the ground pin on the animal’s head. A custom-made cassette guard was attached to the cassette using Kwik Cast adhesive (World Precision Instruments) to minimise the gaps between the cassette and the dental cement. Any remaining gaps were filled with KwikSil (World Precision Instruments) to ensure that the probe was appropriately protected. The entire procedure took ∼4 hours. Once it was complete, the animal recovered in its homecage for a minimum of 3 days, with metacam jelly given daily for analgesia.

#### Burgess and Barry

Mice were habituated to handling in the days leading up to surgery, as well as pre-exposed to non-medicated jelly in their home cage. Mice were at least 12 weeks and at least 27g at the time of surgery. On the day of surgery, subcutaneous analgesics were administered (carprieve 5mg/kg). General anaesthesia was achieved with 3% isoflurane in an induction chamber. The head was shaved, and the eyes were lubricated and covered with foil to protect them. The mice were then moved onto a stereotaxic frame. Anaesthesia was reduced and maintained to a breathing rate of 1 Hz, and their core temperature was monitored while they were on a heat-pad. After confirming the appropriate level of anaesthesia and the absence of a pedal reflex, the head was cleaned with hibiscrub and covered with a sterile drape.

Using an aseptic technique, the skull was exposed and care was taken that the skull was parallel to the flat surface of the stereotaxic frame. The skull was then cleaned of overlaying tissue and etched to aid the adherence of dental cement later in the protocol. One or two screws were then placed with a silver wire attached to act as a ground.

After marking the target site, a cylindrical barrier was made using fast-curing RelyX resin cement (3M) over the site. A custom-made head-plate was added over the frontal plates and cemented in place using dental cement (C&B SuperBond). The cylindrical barrier was used to avoid dental cement flowing over the targeted site. Within the cylindrical barrier, a 1mm (or 0.75mm for single shank probes) craniotomy was then performed over the targeted site. Coordinates relative to the bregma: AP 2.0, ML 2.0, DV 3.4, targeting the hippocampus; AP 4.4mm, ML 3.6mm, DV 2.6mm targeting MEC, with the probe targeted within 0.1mm of the transverse sinus. Care was taken not to damage the sinus. Parts of the dura were then removed with a needle. The exposed skull was temporarily covered with a wetted Sugi sponge (Questalpha).

Next, the Neuropixels probe was lowered to the brain surface. The Sugi sponge was removed and the probe was penetrated to the targeted depth using a custom-built motorized stereotaxic arm (adapted from Kopf) set at 6mm/h while sterile saline was used to keep the brain surface wet. Once at depth, the brain was covered with Dura-Gel (Cambridge NeuroTech), the posts were dropped onto the surface of the head (either skull or head plate) and cemented in place. The screws were tightened, the ground connected to the bone screw, and the implant was covered. The entire procedure typically took about 3-4 hours.

The mice recovered single-housed in their home cages until their pre-surgical weight was re-established or at least 48 hours. During recovery, pain relief was given per-orally using carprieve-infused jelly at 5mg/kg.

#### Carandini/Harris

A brief (approximately 1 h) initial surgery was performed under isoflurane (1–3% in O_2_) anesthesia to implant a steel headplate (approximately 15 × 3 × 0.5mm, 1g). In brief, the dorsal surface of the skull was cleared of skin and periosteum. A thin layer of cyanoacrylate (VetBond, World Precision Instruments) was applied to the skull and allowed to dry. Thin layers of UV-curing optical glue (Norland Optical Adhesives #81, Norland Products) were applied and cured until the exposed skull was covered. The head plate was attached to the skull over the interparietal bone with Super-Bond polymer (Super-Bond C&B, Sun Medical). After recovery, mice were treated with carprofen for three days, then acclimated to handling and head-fixation. On one occasion, the following implantation steps were performed directly after the headplate implantation.

Craniotomies were performed on the day of the implantation, under isoflurane (1–3% in O_2_) anesthesia, and after injection of Colvasone and Carprofren. In brief, the UV glue was removed, and the skull cleaned and scored for best adhesion of the cement. The skull was levelled, before performing the craniotomy using a drill. Once exposed, the brain was covered with Dura-Gel (Cambridge Neurotech).

The implantation phase could be performed either while the mouse was awake or during anaesthesia (1-3% in O_2_). During implantation, the implant was held using a 3D-printed cassette holder and positioned using a micromanipulator (Sensapex). After positioning the shank at the surface of the brain, avoiding blood vessels, the probe was inserted at a slow speed (3-5 µm/s). Once the desired depth was reached, the posts were then loosened from the cassette and dropped to contact the skull, before being cemented in place using Super-Bond polymer. Once the cement was dry, a 3D-printed protective casing was added to protect the shank, and affixed to the cassette using a silicon adhesive (Kwik-Sil, World Precision Instruments). Particular care was given to cover any opening. The whole implant was finally wrapped in tape to hold the flex cable of the probe in place.

#### Isogai

All surgical procedures were carried out under anaesthesia of isoflurane (1.5%-2%, 30-80 mins) after an initial induction into anaesthetized state using isoflurane (5%, 1-2 mins). Prior to placing on the stereotaxic frame mouse fur was trimmed using a shaver, the mouse was fixed to a stereotaxic frame, and lubricating eye gel (Lubrithal, Dechra) was applied to the eyes. Body temperature was maintained at 37°C throughout the procedure. Carprofen (5 mg/kg, s.c.) was administered 10 minutes prior to the end of the surgery for postoperative analgesia.

A custom-built titanium head-fixation implant was affixed to the skull using a cyanoacrylate-based adhesive (Histoacryl, Braun Medical) and dental cement (Simplex Rapid, Kemdent and Super-Bond CB, Sun Medical) centred along the lambda on the mouse skull. The headplate was attached after the peritoneum had been removed and the skull and the surface of the skull roughened using a bone scraper (Fine Science Tools). A 1mm ground screw (ThorLabs) was implanted unilaterally anterolaterally to the headplate and fixed in place using a light-cured resin dental cement (3M, Relyx Unicem 2). Once the headplate and ground-screw were secured, the skin of the animal was glued to either the skull or cement, and the remaining exposed area of skull was covered with a removable silicone sealant (Kwik-Cast, World Precision Instrument). The location of craniotomy needed for access to the medial amygdala (AP: −1800; ML: −2300) was marked during this procedure.

On the day of silicon probe implantation, the animal was anaesthetized as previously described, and a 1mm craniotomy was made at coordinates AP: −1800; ML: −2300 with a 0.3mm burr dental drill. For chronic insertions, a 2mm high wall surrounding the craniotomy site was made prior to drilling by layering UV-curable cement (3M, RelyX Unicem 2) around the site, and the craniotomy was performed once this cement was dry.

#### O’Keefe (rats)

Male rats were implanted with either pre-release Neuropixels 1.0 (3A) or pre-release 4-shank Neuropixels 2.0. Rats were anaesthetized with isoflurane in O_2_ (2-3%) in an induction chamber and given pre-operative analgesia subcutaneously (Carprieve 5mg/kg). The head was shaved, and the surgical site was cleaned with Hibiscrub. In later surgeries, rats also received pre-operative subcutaneous injections of dexamethasone (0.7mg/kg) as this improved neuronal yield and stability. Rats were then transferred to a stereotactic frame and covered with a sterile drape. The rest of the surgery took place under aseptic conditions.

The skull was exposed, and connective tissue was carefully removed. When targeting brain areas located underneath the lateral ridge (i.e., amygdala), we carefully detached a small part of the muscle from the lateral ridge and added a small amount of light-curable epoxy (RelyX) to increase skull surface area and provide better protection for the Neuropixels probe. After levelling the skull, we marked out the target site. First, we drilled small holes away from the target site to insert a stainless steel ground screw (typically either in the prefrontal cortex or cerebellum) and additional support screws. Following the insertion of these screws, a small amount of dental cement (C&B SuperBond) was applied to the threads and left to cure for 10min. Next, we performed a craniotomy and durotomy at the target site. Using a Luigs-Neumann micromanipulator or a standard WPI stereotaxis frame, we slowly inserted the Neuropixels probe at maximal depth to target deep-brain regions. Once the probe was inserted, the posts were cemented onto the surface of the skull (C&B Superbond) and left to cure for 10min. The exposed craniotomy was covered first with a thin layer of Dura-Gel (Cambridge NeuroTech) and then with a layer of sterilized Vaseline that covered the small amount of exposed probe (for MB implants, Vaseline was applied to the probe before the surgery). Once cemented in place, the cassette was detached from the stereotactic holder, and the stereotax was removed. At this point the wire extending from the ground screw was connected to the ground cable from the Neuropixels probe and the whole implant was protected with a flexible copper mesh cage. The opening of the copper mesh cage was wrapped with 3M Coban wrap or a surgical tape when the probe was not in use. The entire procedure typically took between 3-5 hours.

Rats were single-housed following surgery and were given analgesic and antibiotic mixed in strawberry jelly for 3 and 5 days post-surgery, respectively (Metacam 1mg/kg, Baytril 1%). Rats treated with dexamethasone received 0.2 mg/kg at day 2 and day 5 post-surgery.

#### O’Keefe (mice)

During the implantation procedure, the probes were secured in the stereotaxic frame, and yaw and pitch axes were adjusted to ensure that the probe shanks were perpendicular to the horizontal plane through bregma and lambda. A hole for the ground screw was drilled in the right frontal plate and five additional screws were distributed around the implant sites (mEC: 4.3mm lateral to the midline, 0.3mm anterior to the sinus; HPC: 2.5mm lateral to the midline, 4mm posterior to bregma) to provide anchoring for dental acrylic. The whole surface of the skull except the implant locations was covered with Super-Bond cement (Sun Medical). Two 2.4 mm craniotomies were drilled over the implant locations and bone was carefully removed. The dura was gently lifted and a small incision was made to facilitate the probe insertion. The probes were slowly lowered (10–20 µm/s) until they reached the target location. The brain was sealed with artificial dura (3-4680, Dow Corning). The probe shanks above the brain surface were covered with Vaseline. The probes were then fixed to the skull using dental acrylic (Simplex Rapid, Kemdent). A copper mesh cage was attached to the acrylic and connected to the skull screw to shield the probe assembly from external noise. The probe’s ground and reference were connected to the skull screw. Recordings were made in external reference mode.

#### Krupic

During the implantation procedure, the probe was secured in the stereotaxic frame. Two holes for the screws and an additional ground screw to provide anchoring for dental acrylic were drilled anterior to the craniotomy coordinates (mEC: 3.4mm lateral to the midline, 0.3-0.5mm anterior to the sinus) and the screws were inserted in the skull. The headplate was fixed on the skull around the screws and the craniotomy site using Super-Bond cement (Sun Medical). The craniotomy was drilled over the implant location and the bone was carefully removed. The dura was gently lifted and a small incision was made to facilitate the probe insertion. The probe was tilted by 5-10 degrees from anterior to posterior in the sagittal plane and moved above the craniotomy coordinates. The probe was slowly lowered until it reached the target location (at a depth of 2.7-3mm from the brain surface). The probe shanks above the brain surface were covered with Vaseline. The probes were then fixed to the skull using dental acrylic. The probe’s ground was soldered to the ground skull screw.

#### Stephenson-Jones

Mice were anaesthetised with isoflurane (0.5–2.5% in oxygen, 1 L/min) - also used to maintain anaesthesia. Carpofen (5 mg/kg) was administered subcutaneously before the procedure. Craniotomies were made using a 1- mm dental drill (Meisinger, HP 310 104 001 001 004). Prior to training, animals underwent an initial surgery where the skull was exposed, coated with a thin layer of dental cement, and marked for later skull levelling. A craniotomy was drilled over an arbitrarily chosen region of posterior cortex and a ground pin was implanted. To replicate the weight and size of the eventual implant, mice were trained on the task with a size and weight-matched dummy implant, this was fixed during initial surgery to the cement layer using silicon (Kwik-sil: World Precision Instruments). For probe implantation, the dummy implant was removed, and the skull was levelled using previously noted skull markings. A craniotomy and durotomy was made at coordinates AP 0.8mm, ML = 2.1mm and the probe was implanted to a depth of 4.0mm at a 10-degree angle (tip tilted away from midline). The external grounding wire was fixed to the previously implanted skull pin. The craniotomy was then sealed using Duragel (Cambridge Neurotech) and a 3D printed casing was fitted around the probe for protection. In some cases, animals were reimplanted with a second probe and at the end of the experiment probes were recovered for future reuse.

#### Uchida

All procedures were performed in accordance with the National Institutes of Health Guide for the Care and Use of Laboratory Animals and approved by the Harvard Animal Care and Use Committee. Mice were anesthetized with isoflurane (inducted with 4%, maintained at 1.5% in oxygen, 1 L/min). Buprenorphine (SR 0.6 mg/kg, intraperitoneal injection) was administered to provide analgesia. The mice were then moved to a stereotaxic frame. Local anesthetic (bupivacaine) was applied to the scalp to cover the incision site and the eyes were lubricated with ointment (Puralube). Body temperature was maintained at 37°C using a feedback-controlled heat pad (Harvard Apparatus).

A custom headplate and ground pin were implanted as previously described (citation). Briefly, the stereotaxic frame was adjusted to ensure bregma and lambda were in the same horizontal level. The skull surface was etched for 30 seconds (Enamel Etchant Gel; C&B-Metabond) before the headplate was affixed by dental cement (C&B-Metabond). A small craniotomy was drilled in the posterior skull, which was used for the insertion of a ground pin that was also affixed by dental cement. A craniotomy approximately 0.6mm in diameter was drilled at the target site (AP +2.5, ML 1.0) and covered by a silicon sealant (KwikCast, World Precision Instruments). Animals were then allowed to recover, receiving analgesia for at least 48 hours after surgery.

The probe insertion was performed at least five days post the initial surgery. Mice were anesthetized with ketamine/dexmedetomidine (60 mg/kg, intraperitoneal injection) and headfixed using the headplate previously implanted. The silicon sealant was removed, and the area cleaned with saline.

The probe was inserted slowly using a micromanipulator (Luigs & Neuman) into the craniotomy until reaching the desired depth (4 mm DV). Artificial dura (Cambridge Neurotech) was applied on the craniotomy around the probe shank. The posts were then loosened from the cassette and dropped to contact the skull. Two cement (C&B-Metabond) layers were applied to fix the post feet to the skull, waiting 15 minutes between layers to allow for setting. Once the second layer dried, a custom-made plastic guard (3D printed, white ASA) was attached around the exposed shank using layers of a silicone adhesive (Kwik-Sil, World Precision Instruments). The contacts of the connector were protected by a ZIF connector (FH26W-45S-0.3SHW, Hirose Electric Co) attached to the cassette. After implantation, the anesthesia was then reversed using atipamezole (Antisedan, 0.5 mg/kg, intraperitoneal injection).

#### Wills and Cacucci

All subjects were lister hooded rats (Charles River, UK and Envigo, UK). Rat pups were implanted at ages P13-P29 (weighing 28.5-83.0g), and adult rats (above P60) were implanted at 297-327g. Subjects were implanted with three types of silicone probes from Neuronexus (Neuronexus, USA). These were 32 channel probes arranged in a single shank or on four shanks with eight contacts on each shank (A1x32-Poly3-5mm-25s-177-CM32; 4x8-Buzsaki32-CM32, Neuronexus, USA), and 64 channel probes composed of two four shank 32 channel arrays (M4x8-5mm-Buz-200, platform spacing 200um, Matrix-MCM64, Neuronexus, USA). The dimensions of the 3D prints were adapted to fit the Neuronexus probes conserving the same mechanism for implantation and extraction (see modified components in Supplemental Materials: Neuronexus). The dimensions of the guard were also modified and printed in aluminum to provide additional protection from littermates and mothers. Ahead of surgery probes were glued onto the prints as described in the provided protocol. The probes were glued overhanging beyond the post feet allowing for the full probe shank or shanks to be implanted if needed. A difference with the provided protocol was that the posts were attached to the cassette at a 1.5mm overhang before surgery and not moved during the surgery. The post feet were maintained in the modified design but the ‘toes’ were removed to allow for maximum space for the flow of cement between the cassette and the post feet.

Rats were given 3% isoflurane over 4 % oxygen for induction into surgery and 0.5-2% to maintain anaesthesia. A subcutaneous injection of meloxicam 1:10 at 0.1ml per 10g of animal was administered as analgesia (Boehringer Ingelheim; Ingelheim, Germany). Implant coordinates were varied to account for different ages at the time of the surgery. The coordinates used were all measured from bregma at AP: 2.60 to 3.20 mm, ML: 1.80 to 2.20 mm, DV: 2.65 to 2.85 mm below implant site. The DV coordinates were chosen by recording electrophysiological activity during the surgery using the Axona DACQ system (Axona Ltd, UK). The craniotomy was made using a trephine drill of 2 mm diameter, after which the dura was fully removed to provide enough space for the probe arrays. In addition to the craniotomy, five skull screws and a ground screw were added to anchor the dental cement and to provide a grounding gold pin for a flying ground attached to the headstage during recording. Once the probe was in place the craniotomy was covered with Kwik-Cast^TM^ (World Precision Instruments). Deviating from the provided protocol, no cement was applied until after the probe was in place. At this stage a few layers of Super-Bond dental cement were added covering the full surface of the skull, the screws and attaching the posts to the skull (Super-Bond, Sun Medical Co. Ltd, Japan). Once this initial layer of cement was dry the guard/case was attached to the probe using Kwik-Sil (World Precision Instruments) and screwed onto the cassette. Another layer of dental cement was used to attach the case to the skull and provide additional support to the set-up where possible without cementing the cassette to the posts or case. Additional spots of Kwik-Sil were applied to the set-up to cover any visible gaps.

### Explantation

Explantation can be performed both awake or anesthetized. If anaesthesia is used, animals were induced in an induction box, then placed on a stereotaxic frame under isoflurane anaesthesia. Animals were secured to the stereotaxic frame via head posts or ear bars. We calibrated the animals’ head to be perpendicular to the z-axis of the stereotaxic arm to ensure that the probe could be pulled straight. The probe cover and headstage holder were removed to expose the upper holes of the probe cassette, and the probe retrieval prong was connected and secured with screws. We then loosened the screws connecting the posts to the probe cassette. The probe cassette was slowly lifted while carefully monitoring any bending or jittery movements. Explantation typically took 30 minutes. The explanted probe was cleaned in a solution of 1% Tergazyme, followed by distilled water, and isopropyl alcohol. The brains were perfused and extracted for histological sectioning and shank trajectory tracing.

### Survey

We reached out by email to each person that Repix had been shared with, 16 in total. 14 of the users responded to our survey. Users were invited to share their experience with Repix in a semi-structured way using a template (Table S2). To increase the response rate, each person was also offered to simply share summary data of the number of procedures attempted and a success-and-failure rate. Finally, reminders were sent twice to each person. The data was presented individually, as well as collated and summarized.

Users were allowed to define success and failures in each of the periods (implants, data collection, explant) as their experimental needs dictated. Examples of failures in each period include broken shanks, ZIF-connectors chewed by cagemates, implants falling off because of insufficient cementing to the rails, and cement spilled onto the cassette.

Analysis throughout is conducted in MATLAB 2023b. Descriptive summary statistics are reported. Non-parametric Wilcoxon rank-sum tests are performed using MATLAB functions *ranksum* or *signrank*, depending on independent or paired samples, respectively.

### Electrophysiology

We analyze the electrophysiology data from seven users (DR, MH, TP, CM, SB, EM, and CB). All data were acquired using the Neuropixels PXIe acquisition card in a PXI chassis as described in Steinmetz et al^3^, and recorded using either SpikeGLX or OpenEphys. Recordings from Neuropixels 3A probes were acquired using an FPGA card (KC705, Xilinx) which sent data to an acquisition computer. Across labs, animals were engaged in either foraging, social behavior, or headfixed navigation tasks. To standardize between users, the yield was defined as output units after processing raw data with Kilosort 2.5, run with standard parameters (as opposed to the yield after manual curation, which adds inter-user variability^35^). The output was total units, good units and their associated amplitudes. These data were collected as part of other, ongoing experiments, so there was no standardization of recording frequency or duration.

As a descriptive function, exponential lines were fit to the yield data across implantation days using MATLAB function *fit*, with the slope of these lines representing the stability of the yield.

#### Akrami lab

The good unit yield data from VP/EM was further curated using the follow steps: Spike clusters identified during the spike sorting process were included if they were labelled as “good” units by the sorting algorithm and if they passed 3 quality control criteria (amplitude > 50 μV; noise cut-off and refractory period violation), inspired by the International Brain Laboratory preprocessing pipeline^36^.

#### O’Keefe lab

The electrophysiology data (from CM) is mainly, but not exclusively, previously published^22^.

#### Carandini/Harris lab

All recordings were performed during head-fixation. During recordings, the ground and reference were shorted, and connected to the headplate of the mouse. Spikes were clustered using pykilosort^37^, a python port for KS 2.0, with drift correction.

### Measurement of head-angles during behaviour

Accelerometer signals were recorded using a custom IMU sensor board (MPU9250) from the Champalimaud Foundation Hardware Platform, Lisbon, Portugal. The IMU was plugged into a Raspberry Pi mircrocomputer which was synchronised with the Neuropixels acquisition system and the video recording system via a shared external TTL trigger. The acquisition was carried out at 50Hz. Raw signals from the IMU were preprocessed using an adapted version of the scripts developed by Arne Meyer and located in the repository: https://github.com/arnefmeyer/IMUReaderPlugin/tree/master/Python. Raw data signals are converted to roll and pitch angles using these scripts and the angles recorded to a file which is then transferred from the Raspberry Pi for further analysis.

## Protocol

A full procedure protocol can be found at https://www.protocols.io/view/protocol-for-repix-reliable-reusable-versatile-chr-davx2e7n

An initial version of this document was shared with all users. Note that users were not obliged to strictly adhere to the protocol and were encouraged to make changes as they saw fit for their use cases.

## Supporting information

CAD files and materials list

## Data and code availability

The data associated with Repix can be found at https://figshare.com/articles/dataset/Horan_et_al_2024_Repix_reliable_reusable_versatile_chronic_Neuropixe ls_implants_using_minimal_components/25663170

The code to produce the figure panels and analysis can be found at https://github.com/mattiashoran/Repix

## Acknowledgement

We thank T. Mrsic-Flogel and M. Carandini for discussion and supervision, R. Hayman for help in surgical training, and L. Teachen and S. Rajagobalan for animal support. Finally, we thank the Sainsbury Wellcome Centre Neurobiological Research Facility and FabLabs for technical support and manufacturing. **Fig 1** contains a panel that used Biorender.

This work was supported via the Sainsbury Wellcome Centre PhD Programme for MH, DR, SB, ET, AL. MH additionally received grants from Reinholdt W. Jorck og Hustrus Fond, Knud Højgaards Fond and Anglo-Danish Society. CB was supported by EMBO (ALTF 740–2019 fellowship). FC was supported by European Research Council (DEVMEM) and BBSRC (BB/R009872/1). TW was supported by Royal Society (RGF\EA\180119) and Wellcome Trust (220886/Z/20/Z). AA was supported by Gatsby Charitable Foundation (GAT3755) and Wellcome Trust (219627/Z/19/Z). MS was supported by a European Research Council grant (Starting #557533). CB funded by Wellcome SRF (212281/Z/18/Z). NB funded by Wellcome Principal Research Fellowship (222457/Z/21/Z). NB, FC and CB funded by Wellcome Collaborative Award (214314/Z/18/Z). JOK was supporte by a Wellcome Trust Principal Research Fellowship (Wt203020/z/16/z), and a Kavli Foundation grant ( LS-2022-GR-25) to CM and JOK. YI was supported by Gatsby Charitable Foundation (GAT3361) and Wellcome Trust (090843/F/09/Z). This work and AA, MS, YI, JOK were supported by the Sainsbury Wellcome Centre Core Grant from the Gatsby Charitable Foundation and Q22 Wellcome Trust (090843/F/09/Z).

## Supplemental figures

**Supplemental Table 1.**
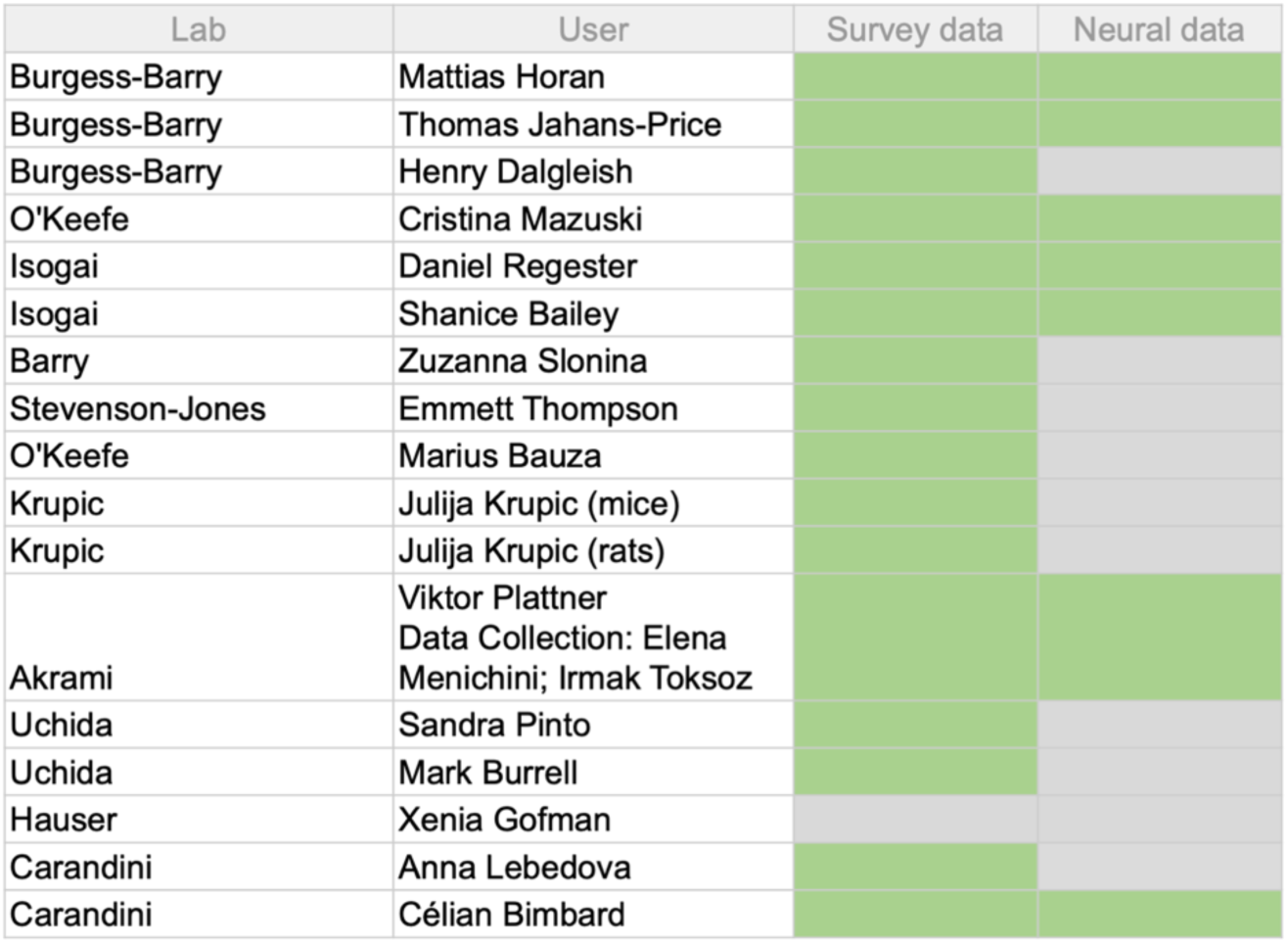
Overview of researchers and laboratories. Repix was shared with sixteen users. Fifteen responded to our census and seven provided electrophysiological yield data (green).

**Supplemental Table 2.** Questionnaire used for the census. Users were sent the above questionnaire to supply details of each procedure attempted and its outcome, along with meta-data.

| ID | Lab | Surgeon | Implant outcome | Data Collection | Explant outcome | Surgery date (implant) | Surgery date (explant) | Implant type(s) | New or reused probe? | Animal | Genotype | Gender | Mount type(s) | Ground screw | Brain region(s) | Coordinates | Durotomy? | Anesthetized or awake | Dexamethasone | Notes |
| --- | --- | --- | --- | --- | --- | --- | --- | --- | --- | --- | --- | --- | --- | --- | --- | --- | --- | --- | --- | --- |
| Animal ID | Primary group | Name | Successful (Y/N)? | Did the probe "survive" data collection? N: e.g. implant came off, shanks broke because of shielding issues. | Successful (Y/N)? | Date | Date | Dummy/Test/Npx3A/Npx1/Npx2 single shank/Npx2 multishank |  | Mouse/rat | Line | M/F | Leg length, if relevant (11mm vs 13mm) | Y/N. Placement, if relevant | Target brain area | Coordinates, including any angles (AP, ML, DV, angles) | Y/N |  | Y/N | Any other notes |

**Supplemental Figure 3.**
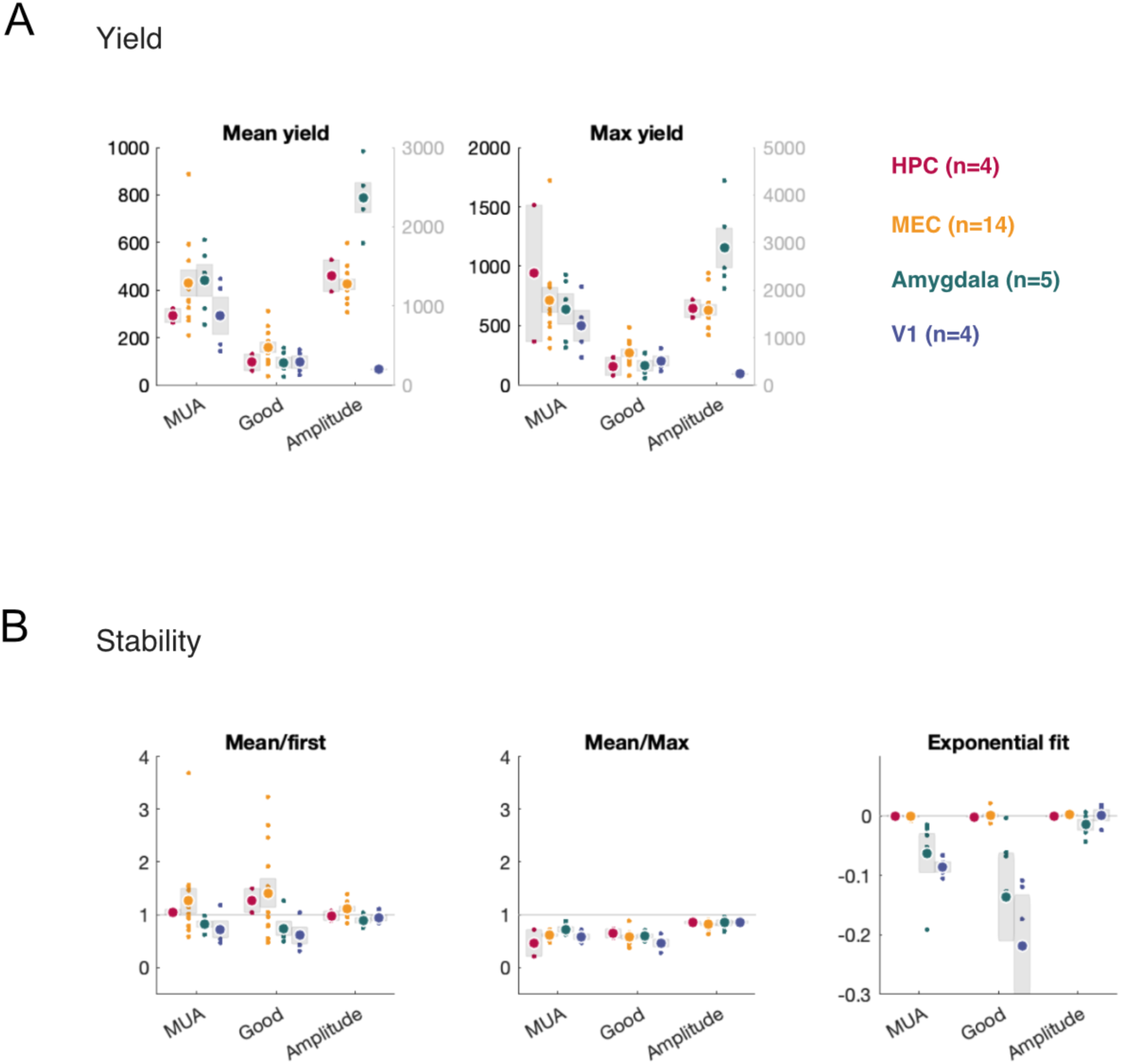
Yield summary. **A:** The mean yield per implantation, left, and max yield per implantation, right. **B:** Three estimates of the stability of the yield: Left, the mean yield divided by the first recording yield, left; middle, the mean yield divided by the max yield at recorded on that implantation; right, the exponential fit to the yield curves over time. Total count, including MUA, only good units, and their associated amplitude, left, middle and right in each panel. Color conventions as in Fig 3.

**Supplemental Figure 4.**
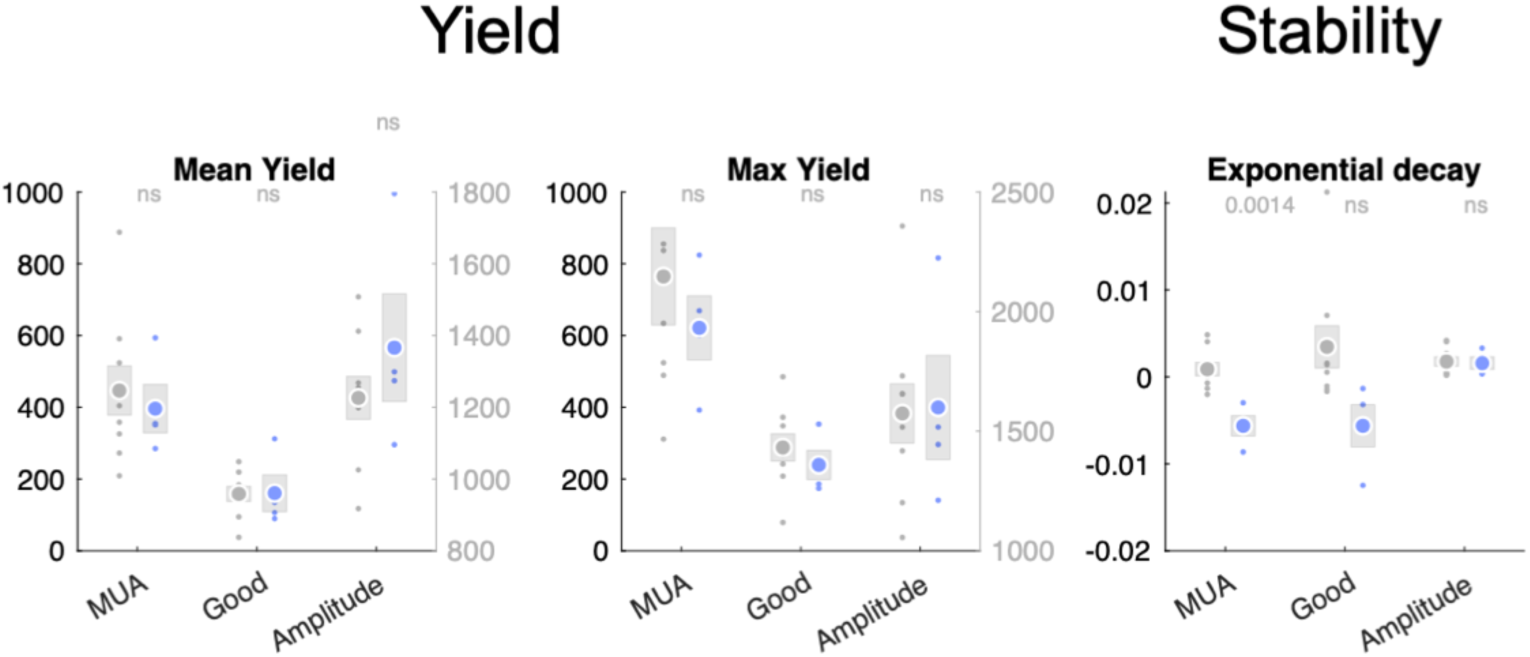
Reuse summary. No significant difference between new (gray) and reused (colored) probes in the entorhinal cortex. The mean yield per implantation, left, and max yield per implantation, middle. The exponential fit to the yield curves shows a significant difference between new and reused probes only in MUA+good units (ie. total units) and a non-significant difference in Good units. Waveform amplitudes are unchanged.

