## Supplementary material for "Repix: reliable, reusable, versatile chronic Neuropixels implants using minimal components": CAD files and materials list: 2-cassetemod_v4_drawing_v1.pdf

**Bottom**

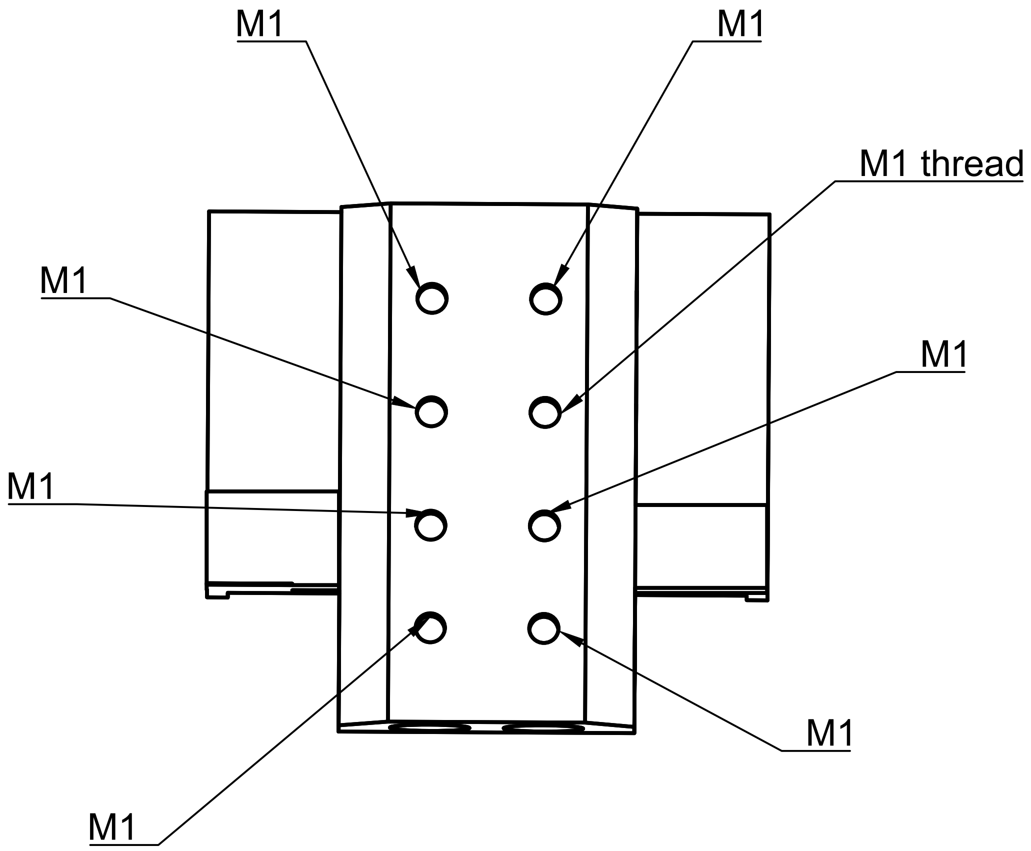


There should be a total of 16 M1 threads on the piece.

8 on the bottom and 4 on each side.

**Right**

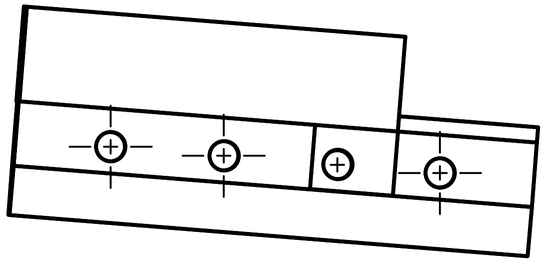

**Left**

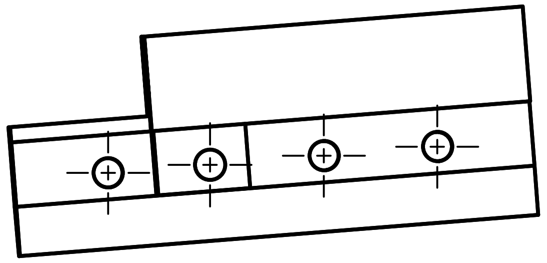

All center marked holes should have an M1 thread.

| Dept. | Technical reference | Created by | Approved by |
|---|---|---|---|
| | | Isabella Varsavsky 18/09/2021 | |
| | | Document type | Document status |
| | | Title<br>CasseteMod_v4 | DWG No. |
| | | | Rev. / Date of issue / Sheet 1/1 |
