## Supplementary material for "Repix: reliable, reusable, versatile chronic Neuropixels implants using minimal components": CAD files and materials list: README.txt

The Repix protocol the for procedure is on protocols.io

____________________________________________________________________________________________________________________________

All .step files are adjustable CAD files i.e. you can open them in Fusion3D or AutoCAD etc. and make adaptations as needed.


CNC Machined (aluminium 6081) parts are:
- cassette 
- posts

3D printed parts are:
- Connector
- Headstage holder
- Cover

____________________________________________________________________________________________________________________________

The PNG file drawings 'cassette_drawing.png' etc. are there for when ordering parts from certain manufacturers which require technical drawings of the parts.

____________________________________________________________________________________________________________________________
