## Supplementary material for "Repix: reliable, reusable, versatile chronic Neuropixels implants using minimal components": CAD files and materials list: Repix_Materials.pdf

#### Repix – core parts

| Part | Material | Versions | File and location |
|---|---|---|---|
| **Cassette** | Aluminium | Standard<br><br>DualProbe prototype | CAD/Cassette/Cassette.step<br><br>CAD/Cassette/Prototypes/DualProbeParallel.step |
| **Posts** | Aluminium | Standard is 11mm in length.<br>Some users have extended this to 13mm. | CAD/ post/post.step |
| **Cover** | PLA | Standard<br><br>Simplified<br><br>"Box and base" Additional protection and miniaturization | CAD/Cover/Standard/Cover.step<br><br>CAD/Cover/Simplified/Cover_simplified.step<br><br>CAD/Cover/BoxBaseCover/Cover_BoxAndBase.step |
| **Headstage holder** | PLA | Neuropixels 1.0<br><br>Neuropixels 2.0 with cover | CAD/HeadStage Holder/headstage_holder_npx1.step<br><br>CAD/HeadStage Holder/headstage_holder_cover_20.step |
| **Connector** | PLA | Standard<br><br>Some user preferred a thicker version which requires sanding. There is also an 8mm slot. | CAD/Connector/Connector.step<br><br>CAD/Connector/Connector_ThickerPlugs.stl<br>CAD/Connector/Connector_8mm.stl |
| **Stereotaxic rod** | Stainless steel. Attach connector to this. | 6mm is standard. Though if a different diameter fits your stereotax, it is easy to change the diameter of the connector-slot to match those needs, e.g. the Connector_8mm.stl fits an 8mm stereotaxic rod. | |
| **M1 screws** | Stainless steel, M1x3 | | |

#### Repix – tools

| Part | Use | Supplier | Alternatives |
|---|---|---|---|
| **Probe holder** | Hold the arm while applying the probe to cassette | Fisso | Noga |
| **Box with graph paper x2** | Align probe | | |
| **Light source** | Cast a shadow from the probe and connector to the graph paper to check alignment of shanks of the probe | Any flashlight, e.g. use your phone. | |

| | | |
|---|---|---|
| **Tweezers** | Curved is easiest for holding the M1 screws | Fine Science Tools (e.g. 91117-10) |
| **Soldering iron** | Short the GRN and REF on the probe. Solder to the ground screw. | |
| **Waveform generator** | Optional, for testing probe after explantation | |

Repix – consumables

| Part | Use | Supplier | Alternatives |
|---|---|---|---|
| **Blu-Tack** | Temporarily hold and cushion probe PCB to the cassette, before applying epoxy | Bostik | |
| **Rapid Epoxy Syringe 24ml** | Epoxy probe to cassette | Araldite | Gorilla Epoxy 25 ml |
| **Weighing boats** | For mixing epoxy | | |
| **Blunt needle and syringe** | For applying epoxy, e.g. 30G | | |
| **DiI** | Fluorescence for shank track tracing, if using | 1 mM in isopropanol, Invitrogen | |
| **Vaseline** | Ensure posts don't stick on explantation | | |
| **Sandpaper** | If needed to sand connector teeth, depending on exact 3D print and how snug the connector sits in the holes of the cassette. | | |
| **Silver or stainless steel wire** | For grounding and e.g. for shorting the GRN and REF: optional depending on experimental needs. | World precision instruments | |
| **Headplate** | Optional | | |

| | | |
|---|---|---|
| **Tergazyme** | 1%, see standard protocol for cleaning Neuropixels | Alconox |
| **Dowsil cleaning solvent** | For cleaning probe, optional | DOWSIL<br>DS-2025 Silicone Cleaning Solvent |
| **PBS** | For cleaning probe | |

### Surgery – tools

| **Part** | **Use** | **Supplier** | **Alternatives** |
|---|---|---|---|
| **Scissors** | Some users prefer making their incision with scissors | Fine Science Tools | |
| **Scalpel** | Making incisions | | |
| **Bone scraper** | Roughen skull for cementing | Fine Science Tools | |
| **Stereotaxic frame, incl arms** | Precision targeting | Kopf Instruments | |
| **Micro manipulator** | Slowly move the probe into the brain | Sensapex | Luigs-Neumann micromanipulator |
| **Thermometer, heating pad, and other measures for the anesthesia** | Maintaining body temperature | Harvard Apparatus | |

Surgery – consumables

| **Part** | **Use** | **Supplier** | **Alternatives** |
|---|---|---|---|
| **Drugs** | In accordance with local guidelines and rules, for anaesthesia, analgesia. Consider dexamethasone as anti-inflammatory. | | |
| **Hibiscrub** | Sterilization | | |

| | | | |
|---|---|---|---|
| **Puralube** | Eye lubricant | | |
| **Phosphoric acid gel** | Etching the skull, optional | Enamel Etchant Gel | |
| **Ground screw** | 1mm ground screw | ThorLabs | |
| **Drill and drill bits** | e.g. 1-mm dental drill, or alterntives, up to users | Meisinger, HP 310 104 001 001 004 | |
| **Sugi tips, Q tips** | Absorbent | Questalpha | |
| **Sterile saline and pipette** | Irrigate the craniotomy during the prolonged lowering of the probe | | |
| **Cyanoacrylate** | Skin closure | VetBond, World Precision Instruments | Histoacryl, Braun Medical |
| **Fibrolar** | Optional, if risk of bleeding | | |
| **Dura-Gel** | Artificial dura | Cambridge NeuroTech | 3-4680, Dow Corning |
| **SuperBond** | Cementing headplate and feed | C&B SuperBond | Sun Medical |
| **RelyX** | Cementing feet and building craniotomy well. | | Norland Optical Adhesives #81, Norland Products |
| **Dental acrylic** | Cementing feet, alternative to RelyX | Simplex Rapid, Kemdent | |
| **UV gun** | Curing UV cement, e.g. RelyX | | |

| | | | |
|---|---|---|---|
| **Kwik-cast or Kwik-sil** | | World Precision Instruments | |
| **Coban band** | Wrapping implant | 3M | Surgical tape |
