## Supplementary figures and images for "Repix: reliable, reusable, versatile chronic Neuropixels implants using minimal components"

### cassette_drawing.PNG

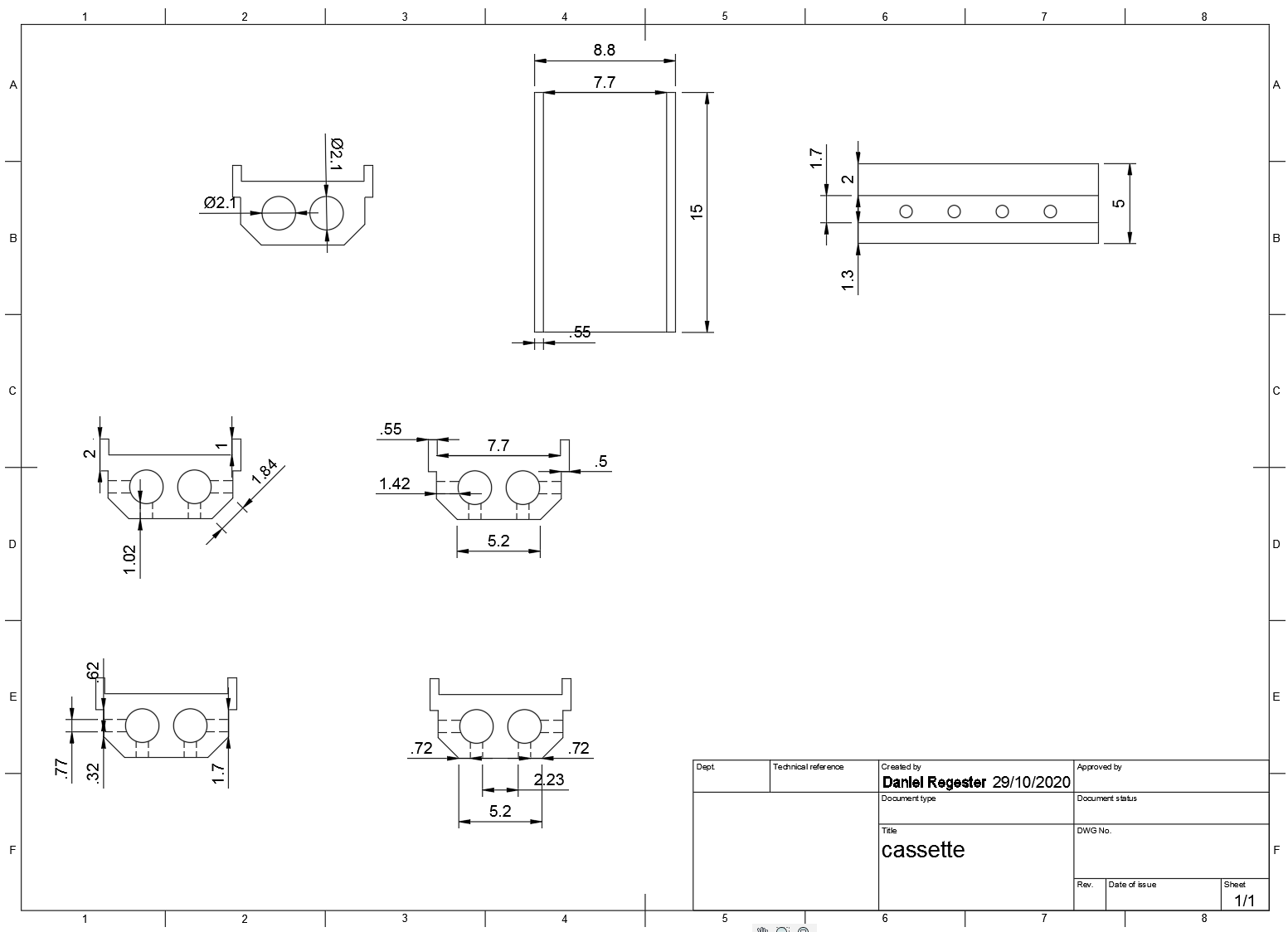

### DualProbe1.png

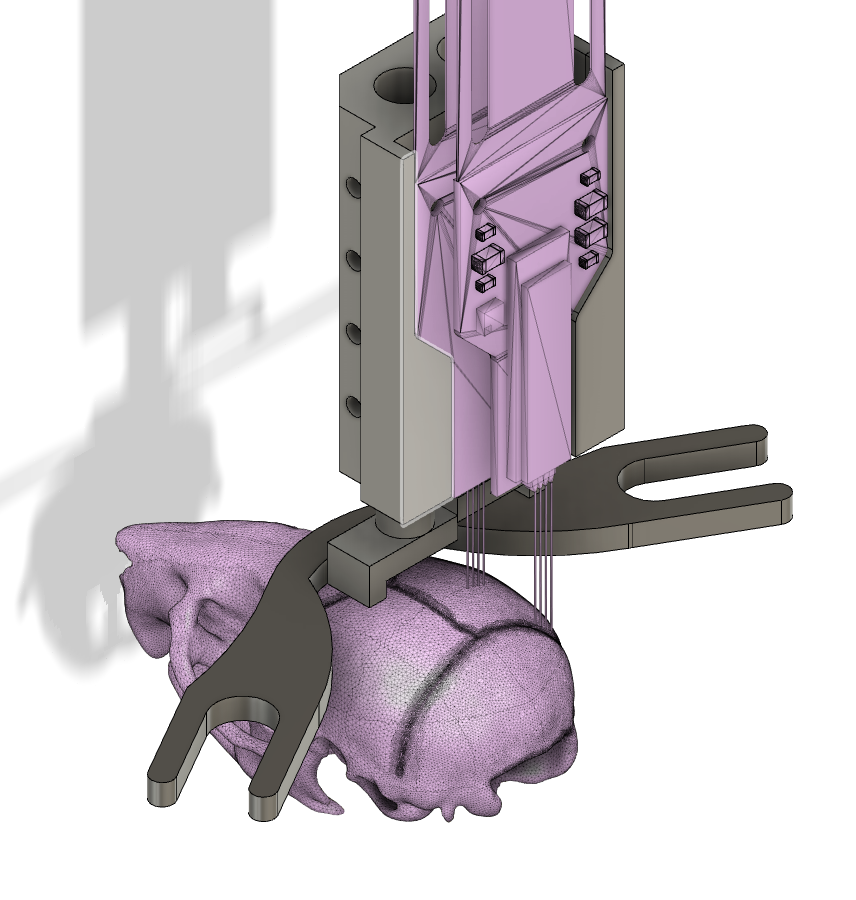

### DualProbe2.png

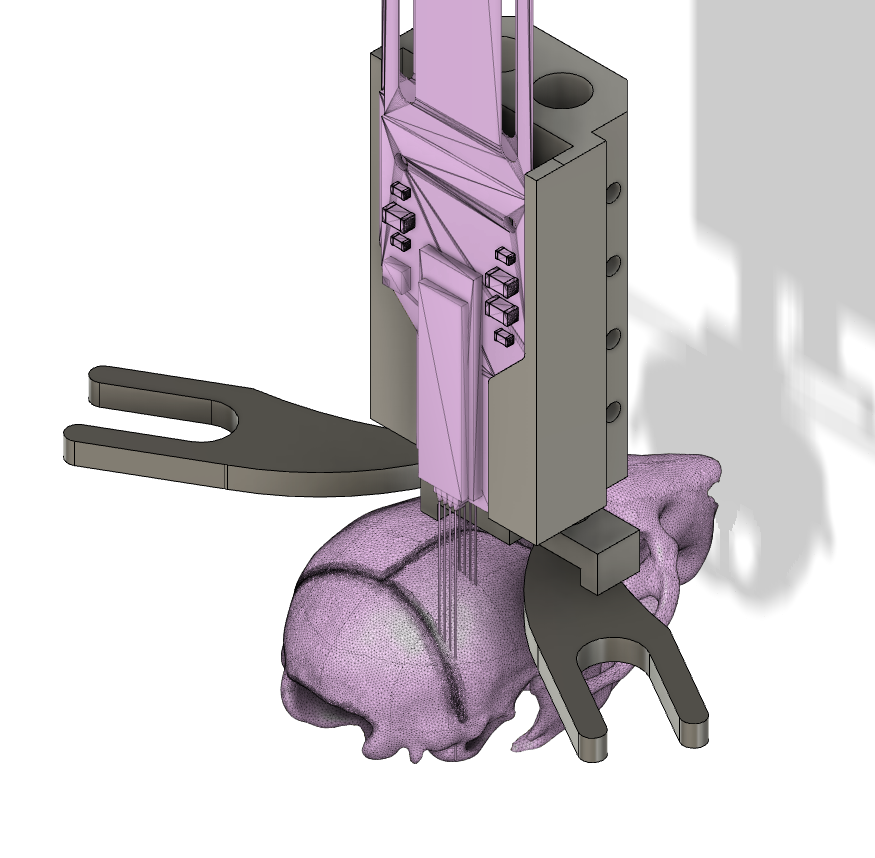

### Post.PNG

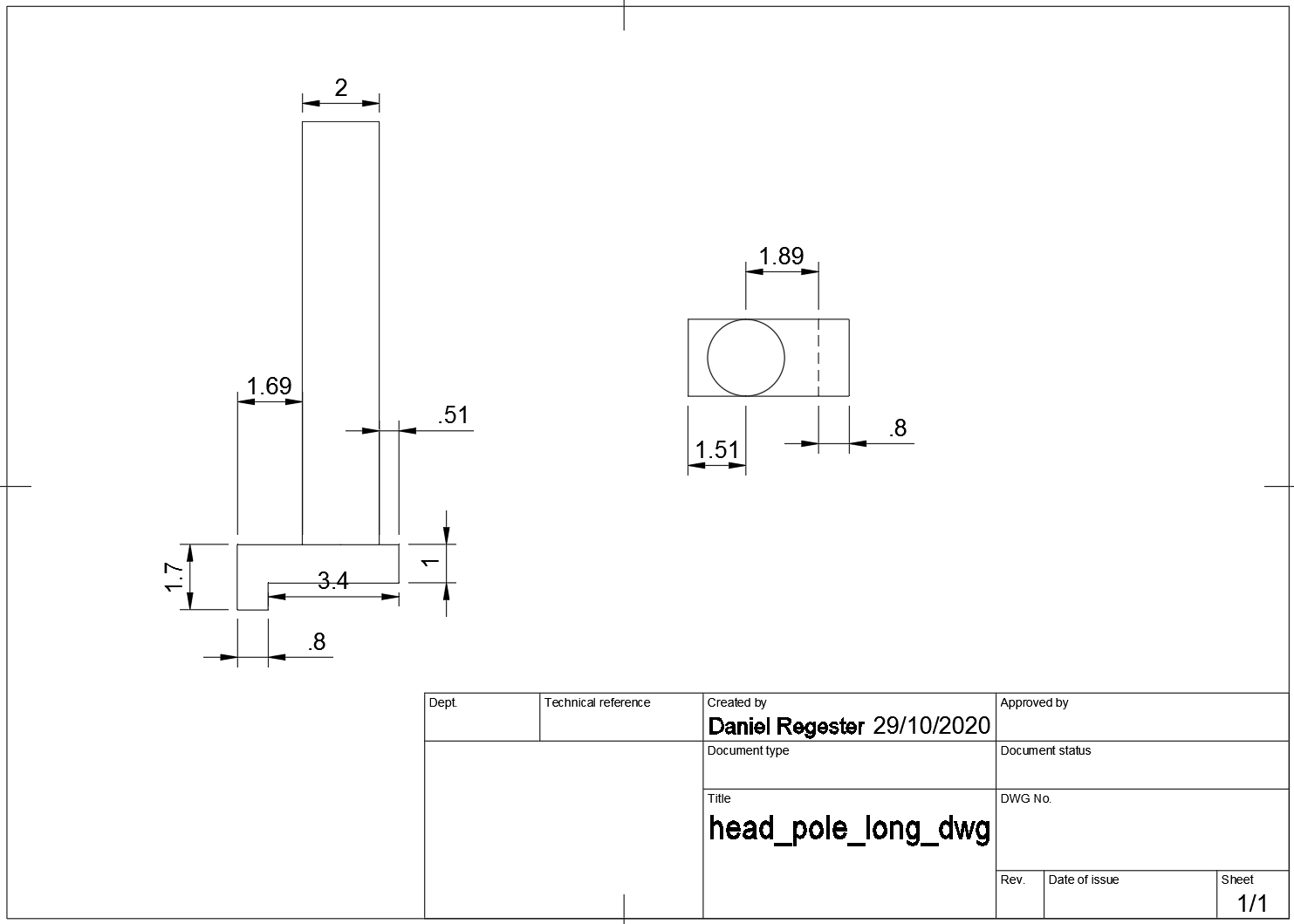
